# Ribonucleoprotein (RNP) condensates modulate survival in response to Mars-like stress conditions

**DOI:** 10.1101/2025.07.09.663920

**Authors:** Riya Dhage, Arijit Roy, Bhalamurugan Sivaraman, Purusharth I Rajyaguru

## Abstract

Scientific advances have emboldened human efforts toward exploring the potential of extinct, extant, or future life on Mars. An important aspect of this endeavor is understanding how an organism adapts to stress-inducing environmental conditions on Mars, such as radiation, shock waves, extreme temperatures, and chaotropic stress due to higher levels of perchlorates. A conserved approach used by organisms across evolutionary scales to adapt and overcome stress conditions is the assembly of ribonucleoprotein (RNP) condensates. In this study, we employ a multidisciplinary approach to understand yeast survivability and adaptation under Mars-like stress conditions, specifically shock waves and perchlorate, by focusing on RNP condensates. Our study reveals that yeast survives 5.6 M intensity shock waves. Exposure to either shock waves or sodium perchlorate induces the formation of P-bodies, a conserved stress-induced condensate. Yeast mutants defective in P-body assembly show defective growth in response to perchlorate stress. Transcriptome analysis, followed by validation, identified several relevant transcripts whose levels are perturbed in response to Mars-like conditions. Finally, identification of several transcripts whose abundance is altered in the P-body assembly mutant upon stress highlights a new connection between response to Martian stress conditions and RNP condensates. This study, a first of its kind, highlights the importance of RNP condensates in understanding the impact of Martian conditions on life in general. This study paves the way for using RNP condensates as a biomarker for assessing the health of life forms during space explorations.

## Introduction

With advances in space science and astrobiology, exploring the potential of Mars in supporting life forms is gaining significant attention. Mars offers a range of hostile environmental conditions that a potential life form would need to overcome. Thus, understanding its unique and challenging environmental conditions becomes important. Martian stress conditions are characterized by the following: a) high-intensity shock waves resulting from meteorite impacts, b) extreme temperature and pressure fluctuations, c) ionizing and solar UV radiations due to a thin atmosphere, and d) chaotropic agents like perchlorates. These conditions pose a serious obstacle to the survival of potential life forms (Beblo-Vranesevic et al., 2020; Cockell & Andrady, 1999). To explore the possibility of extinct/extant/future life forms on Mars, it would be imperative to assess the response of life forms on Earth to Martian conditions.

The Martian land is marked by a history of several meteorite impacts, generating high-intensity shock waves that resonate through the thin Martian atmosphere (Osinski et al., 2020). Shock waves are characterized by rapid changes in pressure and temperature, posing a unique challenge to any potential life forms, as they can induce physiological stresses demanding robust adaptive mechanisms. Exploring the effects of these shock waves on simple eukaryotic microorganisms such as yeast (*Saccharomyces cerevisiae*) could be a good starting point to understand the possibility of life on Mars.

Apart from shockwaves, the presence of perchlorates in Martian soil poses a significant hazard to potential life forms. The presence of perchlorates in Martian regolith has been detected in significant concentrations (0.5 – 1 wt%) at multiple sites, such as the Phoenix landing site (Hecht et al., 2009). Perchlorates are highly oxidizing salts which are chaotropic (disrupting) macromolecules that destabilize hydrogen bonds and hydrophobic interactions. This poses severe chemical stress, making it difficult for life to sustain itself.

Bacteria and yeast have been extensively used as models to study the impact of space and other planetary conditions on life (Bijlani et al., 2021). *Saccharomyces cerevisiae* (a single-celled organism), also known as budding yeast, is a simple yet elegant model system with evolutionarily conserved cellular processes. Its well-documented responses to various stressors make it an ideal model organism for studying the impact of Martian stress conditions on survival. The effect of space travel-related stresses, such as microgravity and ionizing radiation, has been assessed on yeast physiology (Fukuda et al., 2021; Horne et al., 2022; Nemoto et al., 2019; Santa Maria et al., 2023). Specifically, a biosensor using *S. cerevisiae* (BioSentinel) was recently developed by NASA to support deep space missions (Santa Maria *et al*., 2023) to Mars and space habitation beyond low Earth orbit (LEO). These observations highlight the importance of *S. cerevisiae* as a model system for space biology experiments. However, the role of RNP condensates in response to any Martian stresses in altering yeast physiology is currently unexplored.

Assembly of ribonucleoprotein (RNP) condensates like stress granules and P-bodies is a stress response mechanism conserved from yeast to humans. RNP condensates are dynamic, membrane-less structures formed by reversible assembly of RNA and proteins (Buchan, 2014; Buchan et al., 2008). These structures regulate stress response by orchestrating mRNA fate decisions at the level of mRNA stability and translation to control protein abundance. RNP condensates assemble in response to various stress conditions, such as temperature shock, nutrient deprivation, and oxidative stress, and disassemble upon recovery from stress (Buchan et al., 2008; Hoyle et al., 2007). Whether Martian conditions, such as shock waves and perchlorate, induce the assembly of RNP condensates is not known. Additionally, the role of RNP condensates in mounting a response to perchlorate or shock waves alone or in combination has not been explored.

In this study, we employ a multidisciplinary approach comprising engineering, imaging, genetics, and genomics to explore the survivability of *S. cerevisiae* under Mars-like stress conditions, specifically focusing on shock waves and perchlorate stress. We aim to assess whether these stresses could induce RNP condensate assembly and their contribution to yeast survival in these stressful conditions.

## Materials and Methods

### Yeast strains and growth conditions

The yeast strains used in this study are listed in Supplementary Table 1. All strains were transformed with a plasmid expressing Dcp2-mCherry and Pab1-GFP (a list of plasmids used in the study is listed in Supplementary Table 2). All strains were grown at 30°C in synthetic dropout media without uracil supplemented with 2% glucose.

### Shock processing

We utilized the High-Intensity Shock Tube for Astrochemistry (HISTA) housed in the Physical Research Laboratory (PRL), Ahmedabad, India. It is a 7 m long shock tube with a 2 m long driver section and a 5 m long driven section, and these two sections are separated by an Aluminum diaphragm. The driver sections are filled with high-pressure Helium gas, and the driven sections are usually filled with low-pressure Argon gas. The various factors that influence the shock intensity in the shock tube are the bursting pressure of the driver side, the diaphragm’s groove thickness, and the driven gas pressure. By adjusting these parameters, we can generate different shock intensities according to our requirements. The details of HISTA’s operation principles, procedure, and data collection can be found in the literature by (Roy, Singh, Meka, et al., 2022; Singh et al., 2020)

Before the shock processing, we drop casted different yeast samples on the end flange of the shock tube, followed by closing the shock tube’s driven section using the metal end flange.

The entire driven sections are then pumped up to 10^−2^ mbar order, followed by purging using Argon (Ar) to reduce any chance of contamination from the residual atmospheric gas. The driven section is filled with very low-pressure Ar. The driven section pressure can be varied according to the desired shock Mach number. A shock response curve can be collected using two pressure sensors with a separation distance of 30cm attached to HISTA. After the shock processing the internal pressure of the shock tube becomes very high (can be as high as 18 bar). The high-pressure He and Ar mixture is slowly released further with the help of a leak valve. Once the internal pressure of the shock tube equilibrates with the atmospheric pressure, the end flange can be opened, and shocked yeast samples were collected. For the unshocked control, the yeast samples were similarly drop-casted on the metal flange as done for shock processing and for a similar duration of time before collecting them for growth curve and imaging experiments.

### Growth Curve

Growth curve was performed on yeast cells recovered from the shock tube following the exposure to shock waves, along with unshocked cells. The cells were then cultured in fresh media by dilution to an initial optical density of 0.05 OD_600nm._ The cultures were then incubated at 30°C (180 rpm) and monitored the growth periodically till 40 hours.

The growth curve for the perchlorate-treated condition was obtained by growing yeast in fresh media with dilution to an initial optical density of 0.1 OD_600nm._ The cultures were then split into two parts, one treated with 100 mM of sodium perchlorate and one with vehicle control (milli-Q water). The cultures were then incubated at 30°C (180 rpm) and monitored the growth periodically for 30-40 hours.

For combined stress, yeast cells were first exposed to 5.6 Mach intensity shock waves, followed by dilution to an initial optical density of 0.05 OD_600nm_ in fresh medium supplemented with 100 mM sodium perchlorate. The cultures were incubated at 30°C (180 rpm) and monitored the growth periodically till 60 hours. Synthetic dropout media (−uracil) supplemented with 2% glucose were used for all growth curve experiments. All absorbance measurements were performed at 600nm. The data was analyzed, and the growth curve was then plotted using GraphPad Prism 9.0. Lag time and growth rate were derived from the first derivative of the growth curve, subjected to linear regression with Gaussian fit using GraphPad Prism 9.0. For Figure 1C, lag time was calculated as the time taken to reach 0.25 OD_600nm_.

**Figure 1:**
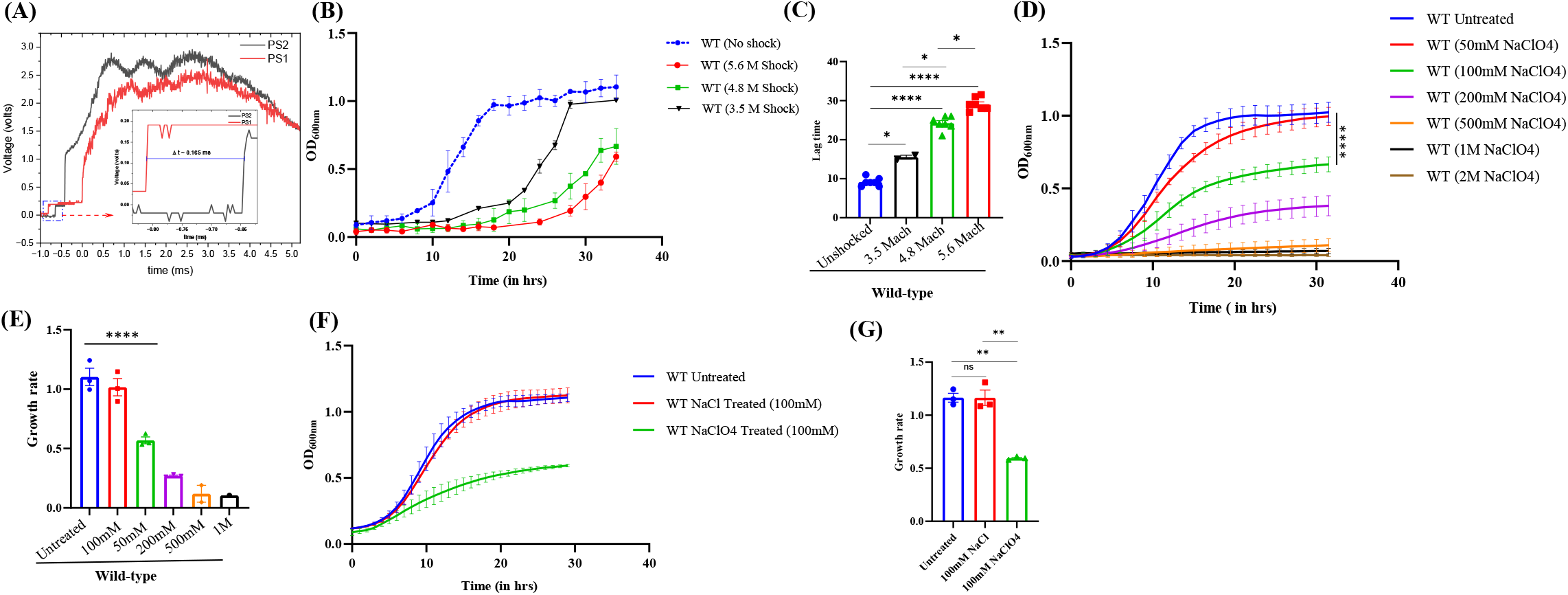
Saccharomyces cerevisiae survives 5.6 Mach intensity shock waves and 100mM sodium perchlorate. [A] Shock response curve of HISTA, used to calculate the Mach value and the reflected shock temperature. [B] Growth curve for wild-type BY4741 yeast strain post-exposure to 3.5, 4.8 and 5.6 Mach intensity shock waves (n=7 biological replicates for 4.8 and 5.6 M, n=2 for 3.5M). [C] Lag time for wildtype BY4741 yeast strain post-exposure to 3.5, 4.8 and 5.6 Mach intensity shock waves obtained from Figure 1B (n=7 biological replicates for 4.8 and 5.6 M, n=2 for 3.5 M). [D] Growth curve for wildtype BY4741 yeast under different concentrations of sodium perchlorate stress (n=3 biological replicates). [E] Growth rate for wildtype under different concentrations of sodium perchlorate obtained from Figure 1D (n=3 biological replicates). [F] Growth curve for wildtype BY4741 yeast under continuous 100mm sodium chloride and sodium perchlorate stress (n=3 biological replicates). [G] Growth rate for wildtype under continuous 100mm sodium chloride and sodium perchlorate stress obtained from 1F. In the above experiments, error bars represent the Standard Error of the Mean. The significance of the data was calculated by Tukey’s multiple comparison test and p-values for all comparisons are summarized in Supplementary Table 4.

### Sample Preparation for Microscopy

Post-shock wave treatment, samples were recovered from the shock tube and fixed with 2% paraformaldehyde in 1X PBS (Phosphate buffer saline pH 7) at room temperature (RT). Fixed samples were then incubated with Poly-L-Lysine-coated coverslips at RT for 45 min, followed by mounting on a glass slide with Fluorochrome G.

For the sodium perchlorate-treated condition, cells were grown in the presence of 100 mM sodium perchlorate to reach 0.5 OD., followed by fixing with 2% paraformaldehyde as mentioned above.

### Microscopy

All images were acquired using a Deltavision RT microscope system running softWoRx 6.1.3 software (Applied Precision, LLC), using a 100X, oil-immersion objective. Exposure time and transmittance settings for the Green Fluorescent Protein (GFP) channel were 0.3s and 32%, and for the mCherry channel were 0.5s and 50%, respectively. Images were acquired at a resolution of 512 × 512 pixels using a CoolSnapHQ camera (Photometrics) with 1×1 binning. All the images were deconvolved using standard softWoRx deconvolution algorithms. ImageJ was used to analyze the data. For each experiment, 100 cells were counted to determine the number of foci (Pab1-GFP, Edc3-mCherry, or Scd6-GFP) per cell. For figure 1F-G, live-cell images were acquired using an Apotome microscope (Zeiss Axio observer) and processed using ImageJ software, as there were technical issues with Deltavision RT microscope system. The images were taken at 63X, oil immersion objective. Exposure time and transmittance settings for the Green Fluorescent Protein (GFP) channel were 0.15s and 50%, and for the mCherry channel were 0.75s and 100%, respectively.

### Colony-forming unit (CFU) assay

The yeast cells were diluted at an OD_600_ of 0.1 from an overnight-grown primary culture and grown in synthetic dropout media without uracil (SD Ura-), supplemented with 2% glucose, with or without 100 mM sodium perchlorate, and allowed to reach 0.5 OD_600_. After which the cultures were washed twice with perchlorate-free media. The cells were serially diluted and plated on SD Ura-agar plates. The plates were incubated at 30 °C for 48 hours. The CFU count was used to determine the survival.

### mRNA stability assay

A thermo-sensitive mutant of RNA polymerase II, *rpb1-1*, was used to perform an mRNA stability assay. (Nonet et al., 1987; Parker et al., 1991). The *rpb1-1* strain was grown in YEPD with or without 100 mM sodium perchlorate at 30°C (180 rpm) till 0.5 OD_600nm_. For transcription shut off, pre-warmed (55°C) YEPD was added to cultures at 30°C followed by incubating at 42°C for heat shock. Cells were collected at time intervals of 0, 15, 30, 45, and 60 minutes of heat shock, followed by immediate snap freezing with liquid nitrogen. RNA isolation was performed by the hot-phenol method, followed by cDNA synthesis. Gene-specific primers were used for qRT-PCR and were normalized to PGK1. Statistical analysis was done by 2-way ANOVA using Šídák’s multiple comparisons test.

### RNA isolation

RNA was isolated from biological replicates for each test sample using the hot acidic phenol protocol described in Current Protocols in Molecular Biology (1996), Unit 13.12 (Collart & Oliviero, 2001). Briefly, the cell pellet was washed with DEPC-treated autoclaved Milli-Q water (DEPC-AMQ) once and resuspended thoroughly in TES (10 mM Tris-Cl pH 7.5, 10 mM EDTA, and 0.5% SDS). 400μl acidic phenol pH 4.5 (0.1 M citrate buffer saturated) was added to the suspension and incubated at 65 °C for 60 min with occasional vortexing. Samples were spun at 14,000 rpm for 10 min at 4 °C. The top aqueous layer was transferred to a fresh tube. Subsequently, 400μl chloroform was added, mixed, and spun as above. The aqueous layer was mixed with 1/10th volume 3 M sodium acetate, pH 5.2, and 2.5 volumes of 100% ethanol, followed by snap freezing in liquid N_2_. The samples were allowed to thaw on ice, followed by pelleting down the precipitated RNA at 14,000 rpm at 4°C for 30 minutes. The RNA pellets were washed with 70% Ethanol (prepared in DEPC-AMQ, followed by air drying and resuspending the pellet in 50μl DEPC-AMQ, and RNA quality was determined by 1% agarose formamide gel electrophoresis. The concentration and purity of RNA were quantified using a Nanodrop Spectrophotometer (Thermo Scientific; 2000). The integrity of RNA in the samples was assessed on TapeStation (Agilent). RNA concentration was quantified using a Qubit RNA HS assay kit (Q32855).

### RNA library preparation and sequencing

RNA sequencing libraries were prepared with Illumina-compatible NEBNext® Ultra™ II Directional RNA Library Prep Kit (New England BioLabs, MA, USA) at Genotypic Technology Pvt. Ltd., Bangalore, India. 200 ng of total RNA was used for mRNA isolation, fragmentation, and priming. Fragmented and primed mRNA was then subjected to first-strand synthesis followed by second-strand synthesis. The double-stranded cDNA was purified using NEBNext sample purification beads. Purified cDNA was end-repaired, adenylated, and ligated to Illumina adapters as per NEBNext® Ultra™ II Directional RNA Library Prep protocol, followed by second-strand excision using the USER enzyme at 37°C for 15 minutes. Adapter-ligated cDNA was purified using NEBNext beads and was subjected to 12 cycles for Indexing-(98°C for 30 sec, cycling (98°C for 10 sec, 65°C for 75 sec) and 65°C for 5 min) and enriched the adapter-ligated fragments. Final PCR products (sequencing library) were purified with NEBNext beads, followed by a library quality control check. Illumina-compatible sequencing libraries were quantified by Qubit fluorometer (Thermo Fisher Scientific, MA, USA), and fragment size distribution was analyzed on Agilent 4150 TapeStation. The libraries were paired-end sequenced on an Illumina NovaSeqX Plus sequencer for 150 cycles (Illumina, San Diego, USA) following the manufacturer’s instructions.

### Transcriptome analysis

The QC-passed reads were mapped to the indexed *Saccharomyces cerevisiae* reference genome (S288c) using a splice-aware aligner. (Langmead et al., 2019). The quantification of expressed transcripts was performed using the genome-aligned reads. This expression data was subsequently utilized for downstream analyses, including gene ontology and pathway annotations, to study the function of each expressed transcript. The Featurecounts4 tool was used to estimate and calculate transcript abundance. (Liao et al., 2014). DESeq25 was used to calculate the differentially expressed transcripts. (Love et al., 2014). Transcripts were functionally annotated based on homology using the BLAST5 tool against Fungal protein sequences available from the UniProt database. (Buchfink et al., 2015). Pathway analysis was done on the KAAS6 server using *Saccharomyces cerevisiae* as the reference organism. (Kanehisa et al., 2021).

### Reverse transcription and quantitative real-time PCR

The isolated RNA was subjected to DNase treatment to remove any DNA contamination. For this, 5 μg of total RNA, 0.75 units of DNase I (Thermo Fisher Scientific EN0525), and DNase I buffer (with MgCl2 added to it), 0.2 units of RiboLock RNase Inhibitor Thermo Fisher Scientific, EO0381), and DEPC-AMQ were added for a 20 μl reaction. The above reaction was incubated at 37°C for 30 minutes, followed by adding 2 μl EDTA (Ethylene-diamine-tetraacetic acid) and incubating it at 65°C for 10 minutes to inhibit the DNase I activity. The DNaseI-treated RNA concentration was measured in a spectrophotometer, followed by an RNA quality check by 1% agarose formamide gel electrophoresis. DNase I-treated RNA was used for cDNA preparation. One μg of RNA was used to synthesize cDNA using the RevertAid RT Reverse Transcription Kit (Thermo K1691) according to the manufacturer’s protocol. The synthesized cDNA was then diluted 1:10, followed by quantitative real-time PCR.

TB GreenTM Premix Ex TaqTM (TakaRa). The reactions were set up in technical duplicates for each biological replicate. Data points in all graphs represent biological replicates. Primers used for qRT-PCR are listed in Supplementary Table 3. The PCR conditions were 95°C for 10 minutes for initial denaturation, followed by 30 cycles of 95°C for 15 seconds and 60°C for 45 seconds. DNA was quantified in every cycle at the step of extension. Melt cure acquisition was carried out at 60°C for 15 seconds. PGK1 was used as an internal control. To calculate the final log2 fold-change values, ΔΔCt method was used, where log2 Fold-change values were plotted using GraphPad Prism software, version 9.00. The significance of the data was calculated by a paired *t*-test.

## Results

### S. *cerevisiae* survives 5.6 Mach intensity shock waves

Using a unicellular eukaryote, *Saccharomyces cerevisiae* (budding yeast) as a model system, our study aims to understand the impact of Mars-like stress conditions on survival and growth. To address this question, we employed a multidisciplinary approach to simulate conditions similar to those on Mars, specifically shock waves. Using a High-Intensity Shock Tube for Astrochemistry (HISTA), the *S. cerevisiae* wild-type BY4741 strain was subjected to varying intensities of shock waves such as 3.5, 4.8, and 5.6 Mach (M) (detailed in the Methods section). The various operational parameters for shock processing used in this study are outlined in Table 1. The shock response curve (5.6 M) of HISTA (Figure 1A) was recorded on a digital oscilloscope using two piezoelectric pressure sensors, PS1 and PS2, positioned at the end of HISTA. The inset highlights the time delay between the pressure signals from PS1 and PS2 during the arrival of the incident shock. It is notable that shock wave treatment also exposes the cells to very high estimated temperatures, albeit for a very short (2ms) period of time (Table 1; Roy, Singh, Ambresh, et al., 2022). This indicates that shockwaves could expose lifeforms to transient heat shock stress. To assess the effect of shock waves on *S. cerevisiae* survival, growth curve analysis was performed on wild-type BY4741 yeast cells collected post-shock from the shock tube. Interestingly, as shown in Figures 1B and 1C, the *S. cerevisiae* wild-type BY4741 strain survives 5.6 Mach intensity shock waves with an extended lag phase compared to the unshocked controls. 5.6 M shocked wild-type strain began to grow after approximately 20 hours of incubation at 30°C, indicating a prolonged recovery period from the stress induced by shock waves. Cells exposed to 3.5 and 4.8M intensity shockwaves also underwent an extended lag phase, but to a lesser extent than 5.6M intensity shockwaves (Figure 1B and C). This indicates a graded response by yeast cells to increasing intensity of shock waves.

**Table 1.** The details of the shock tube parameters used in the experiment.

| Serial Number | Bursting Pressure (He) (Bar) | Driven gas (Ar) Pressure (Bar) | Mach Number (M) | Estimated Reflected Shock Temperature (K) calculated using Rankine-Hugoniot Jump equations |
| --- | --- | --- | --- | --- |
| 1 | 62.5 | 0.1 | 5.5 | 7300 |
| 2 | 30.3 | 0.1 | 4.8 | 5355 |
| 3 | 27.8 | 0.4 | 3.5 | 2915 |

### S. *cerevisiae* survives sodium perchlorate concentrations equivalent to Martian conditions

To further study the impact of Martian stress conditions on *Saccharomyces cerevisiae,* we investigated the effect of perchlorate salts at concentrations equivalent to those found on the Martian surface. The Martian soil is reported to contain perchlorates in the range of 0.5-1% by weight. (Hecht et al., 2009). To test this, we used sodium salt of perchlorate (NaClO4) as it is highly soluble in water among all other perchlorates. We then analyzed the growth of wild-type BY4741 yeast strain in the continuous presence of sodium perchlorate at concentrations of 50 mM, 100 mM, 200 mM, 500 mM, 1M, and 2 M. As shown in Figures 1D and 1E, we observed a concentration-dependent increase in growth defect, indicating the negative effect of sodium perchlorate on yeast growth. While 50mM NaClO4 impacts yeast growth slightly, concentrations of 100mM and 200mM show significant growth defects, and concentrations of 500mM or above result in little or no growth till 40 hours of incubation at 30^0^ C. Since perchlorates are present on Martian soil in the range of 0.5-1%, we have therefore used 100mM (1.22% by weight) of sodium perchlorate for all subsequent experiments.

To check whether the observed growth defect is specific to sodium perchlorate, we looked at yeast growth in the presence of 100 mM sodium chloride. We observed no growth defect for wildtype at 100mM NaCl compared to untreated control, unlike sodium perchlorate (Figure 1F-G). This highlights the specific effect of perchlorate ions on growth.

### 5.6 M intensity shock induces RNP condensates in yeast

Yeast cells survive 5.6M shock waves and grow after an 18-20 hour lag, indicating a stress response. We, therefore, investigated the assembly of two conserved, classical RNP condensates: stress granules (Pab1-GFP) and P-bodies (Dcp2-mCherry) in response to shock waves. Exposure to 5.6 M intensity shock waves induced the assembly of both P-bodies as well as stress granules (Figure 2A). This observation indicates that exposure to shock waves induces the assembly of ribonucleoprotein condensates as a cellular response to this stress. Moreover, the number of stress granules and P-bodies decreased in cells exposed to 4.8M compared to 5.6M (Figure 2B-C). Interestingly, no condensates were induced in response to a 3.5M shockwave, suggesting that the induction of the condensates occurs in a graded manner depending on the shockwave intensity. Since we observed the assembly of P-bodies and stress granules in wild-type strain upon exposure to 5.6M shock waves, we wanted to confirm the assembly of these condensates with another condensate resident marker protein, Scd6, which is reported to localize to both P-bodies and stress granules. As shown in Supplementary Figure 1, the assembly of Scd6-GFP granules was also observed in wild-type yeast strains upon exposure to 5.6M intensity shock. This again confirms our observation that exposure to shock waves induces the assembly of ribonucleoprotein condensates as a cellular response to the stress condition.

**Figure 2:**
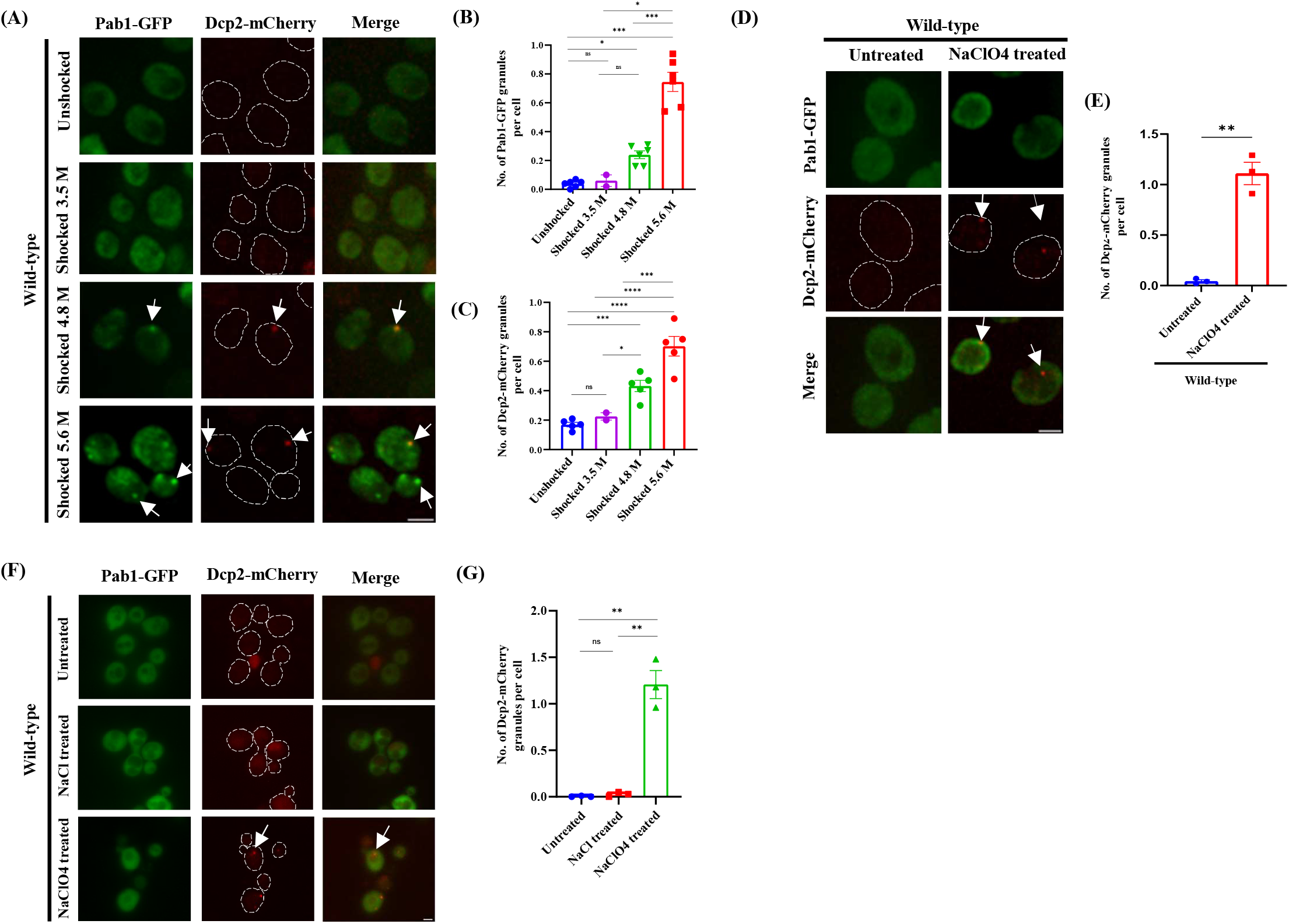
Shock waves and sodium perchlorate induce the assembly of RNP condensates in wildtype (BY4741) yeast strain. Microscopy analysis for wildtype (BY4741) yeast strain - [A] upon exposure to 5.6, 4.8 and 3.5 Mach intensity shock waves, [D] upon continuous 100mM sodium perchlorate treatment,.[F] upon continuous 100mM sodium chloride and sodium perchlorate treatment. Pab1-GFP and Dcp2-mCherry are the markers of stress granules and P-bodies respectively. Arrows in Dcp2-mCherry and Pab1-GFP panels represent P-bodies and stress granules, respectively. Scale Bar, 2 µm. [B-C] Graph depicting the number of granules per cell in wild-type upon exposure to 5.6, 4.8 and 3.5 Mach intensity shock wave for [B] Pab1-GFP, [C] Dcp2-mcherry (n=6 biological replicates for 5.6 and 4.8 M, n=2 for 3.5 M); [E,G] Graph depicting the number of Dcp2-mCherry granules per cell in wild-type strain upon [E] continuous sodium perchlorate treatment [G] continuous sodium chloride and sodium perchlorate treatment (n=3 biological replicates). In the above experiments, error bars represent the Standard Error of the Mean. The significance of the data was calculated by a two-tailed paired student t-test and p-values are summarized as: ^****^, p < 0.0001 ^***^, p < 0.005; ^**^, p < 0.01; ^*^, p < 0.05 *.

### Sodium perchlorate induces P-body assembly

*S. cerevisiae* survived 100 mM sodium perchlorate with a significant growth defect. Therefore, we investigated the assembly of RNP condensates in response to sodium perchlorate stress. As depicted in Figures 2D and 2E, P-bodies but not stress granules were induced in wild-type strain grown in the presence of continuous 100 mM NaClO4. This observation indicates that signalling pathways for the assembly of PBs and SGs are distinct in response to sodium perchlorate stress. To determine whether this response is specific to perchlorate ions or a general effect of sodium salts, we assessed assembly of RNP condensates in the presence of sodium chloride. Notably, neither stress granules nor P-bodies were induced in response to sodium chloride, indicating the effect is specific to perchlorate ions (Figure 2F-G).

### *edc3Δlsm4ΔC* mutant fails to assemble RNP condensates in response to shock waves or sodium perchlorate stress

We wanted to investigate the importance of these condensates in the recovery of yeast cells from the two Martian stress conditions mentioned above. To address this, we examined the mutant strain *edc3Δlsm4ΔC*, which is known to be defective in the assembly of P-bodies and stress granules under stresses such as glucose starvation. (Decker et al., 2007). This strain is deleted for EDC3, the gene encoding a conserved P-body marker, and a part of the LSM4 gene encoding the prion-like C-terminal of the Lsm4 protein. We assessed whether this mutant could assemble RNP condensates in response to shock waves and/or perchlorate stress. As shown in Figures 3A-B and Supplementary Figure 2A-C, P-body and stress granule assembly were hampered in the *edc3Δlsm4ΔC* mutant upon exposure to 5.6 M intensity shock wave and sodium perchlorate. This finding suggests that the formation of Dcp2-mCherry marked P-bodies and Pab1-GFP marked stress granules depends on the presence of Edc3 and Lsm4, highlighting similarities between the condensate assembly in various stress conditions.

**Figure 3:**
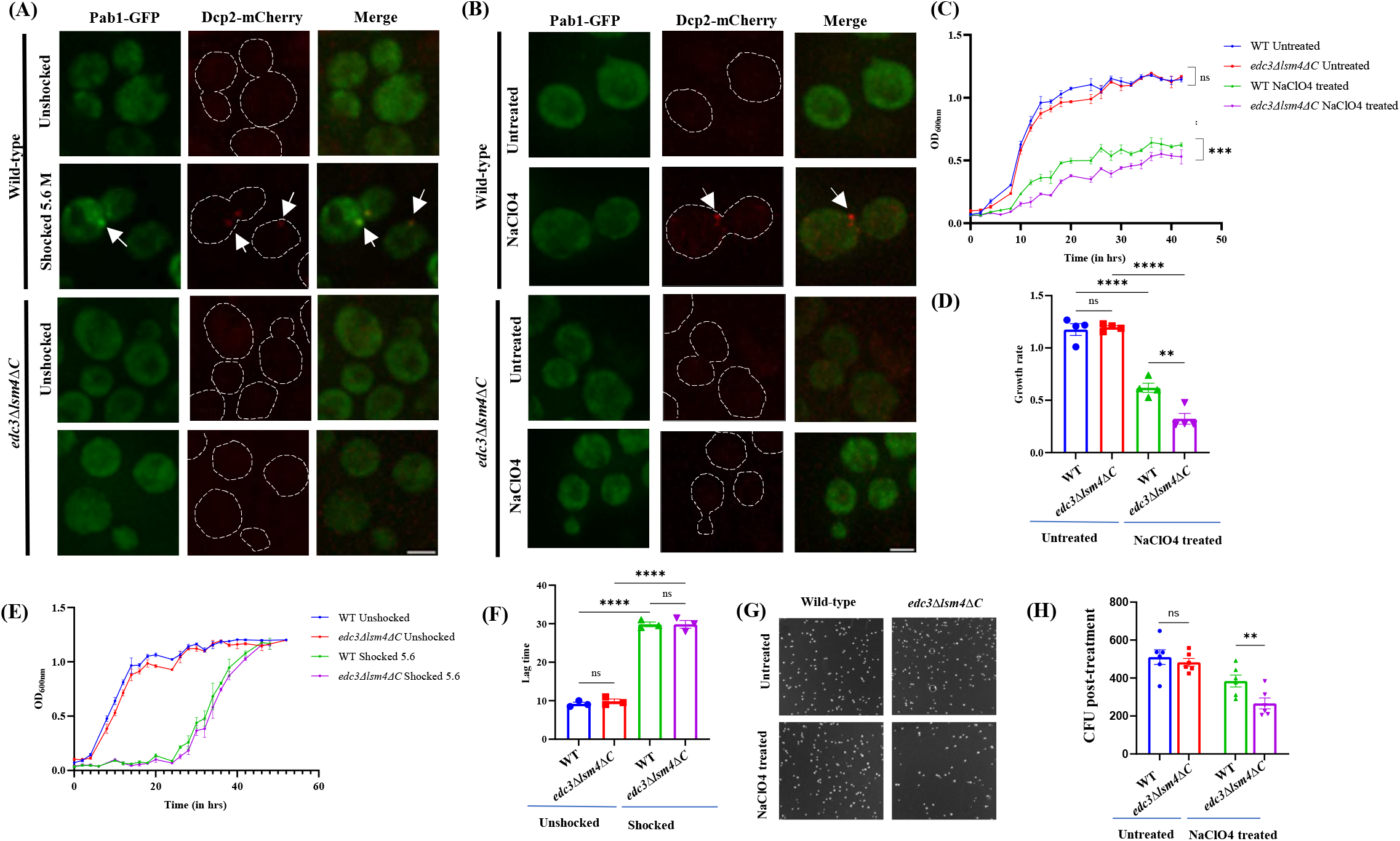
edc3Δlsm4ΔC mutant is defective in condensate assembly and demonstrates an extended lag phase in sodium perchlorate stress. Microscopy analysis for edc3Δlsm4ΔC mutant strain upon - [A] exposure to 5.6 Mach intensity shock waves, [B] continuous sodium perchlorate (NaClO4) treatment. Dcp2-mCherry and Pab1-GFP are the markers of P-bodies and stress granules, respectively. Scale Bar, 2 µm. Comparison of the growth curve for wild-type and edc3Δlsm4ΔC mutant – [C] grown in continuous 100mM sodium, [E] post-exposure to 5.6M intensity shock. [D] Graph depicting growth rate in wild-type and *edc3Δlsm4ΔC* upon sodium perchlorate stress obtained from Figure 3C (n=4); [F] Lag time for wild-type and *edc3Δlsm4ΔC* mutant obtained from Figure 3E (n=3 biological replicates); [G] Survival of wild-type and edc3Δlsm4ΔC post continuous treatment with 100mM sodium perchlorate shown as colony forming units (CFU). [H] Quantification for CFU obtained from Figure 3G (n=6 biological replicates). In the above experiments, error bars represent the Standard Error of the Mean. The significance of the data was calculated by Tukey’s multiple comparison test for growth curves and two-tailed paired student t-test for CFU assay with p-values summarized as : ^****^, p < 0.0001 ^***^, p < 0.005; ^**^, p < 0.01; ^*^, p < 0.05 *.

### The edc3Δlsm4ΔC mutant demonstrates an extended lag phase in sodium perchlorate stress

We hypothesized that if the formation of RNP condensates played a crucial role in survival under these stresses, then the *edc3Δlsm4ΔC* mutant, which fails to assemble these condensates, could exhibit altered growth under these stresses compared to wild-type. Therefore, we assessed the growth of the *edc3Δlsm4ΔC* mutant upon exposure to these stresses. Interestingly, the *edc3Δlsm4ΔC* mutant exhibited significantly slower growth than the wild-type strain (Figure 3C and 3D) in continuous perchlorate stress. This suggests that RNP condensate assembly could facilitate better cellular adaptation and growth under perchlorate stress. In contrast, there was no significant difference in growth between wild-type and mutant strains during recovery after exposure to shock waves (Figures 3E and 3F). Next, we examined the effect of sodium perchlorate on the survival of *edc3Δlsm4ΔC.* To test this, we performed a colony-forming unit (CFU) assay. Our results revealed a significant decrease in CFU count in *edc3Δlsm4ΔC* compared to the wild-type following sodium perchlorate treatment, as shown in Figures 3G and 3H, indicating decreased survival of *edc3Δlsm4ΔC* under these conditions. This is consistent with our earlier observation of slower growth in *edc3Δlsm4ΔC* compared to wild-type during continuous perchlorate treatment.

### Transcriptome analysis reveals that granule assembly affects mRNA abundance in wild-type and *edc3Δlsm4ΔC* in response to sodium perchlorate

RNP condensates regulate stress response by altering the mRNA abundance and translation. To understand the transcriptome-wide changes induced by the perchlorate stress and the possible contribution of RNP condensates to it, we performed sequencing of total RNA samples from wild type and the *edc3Δlsm4ΔC* mutant (Figure 4). For differential gene expression analysis, the category listed first in all comparisons serves as a control. Genes with *p*-value ≤.05 were considered significantly differentially expressed. Figure 4A-C and Supplementary Figure 3 show a summary of differentially expressed genes.

**Figure 4:**
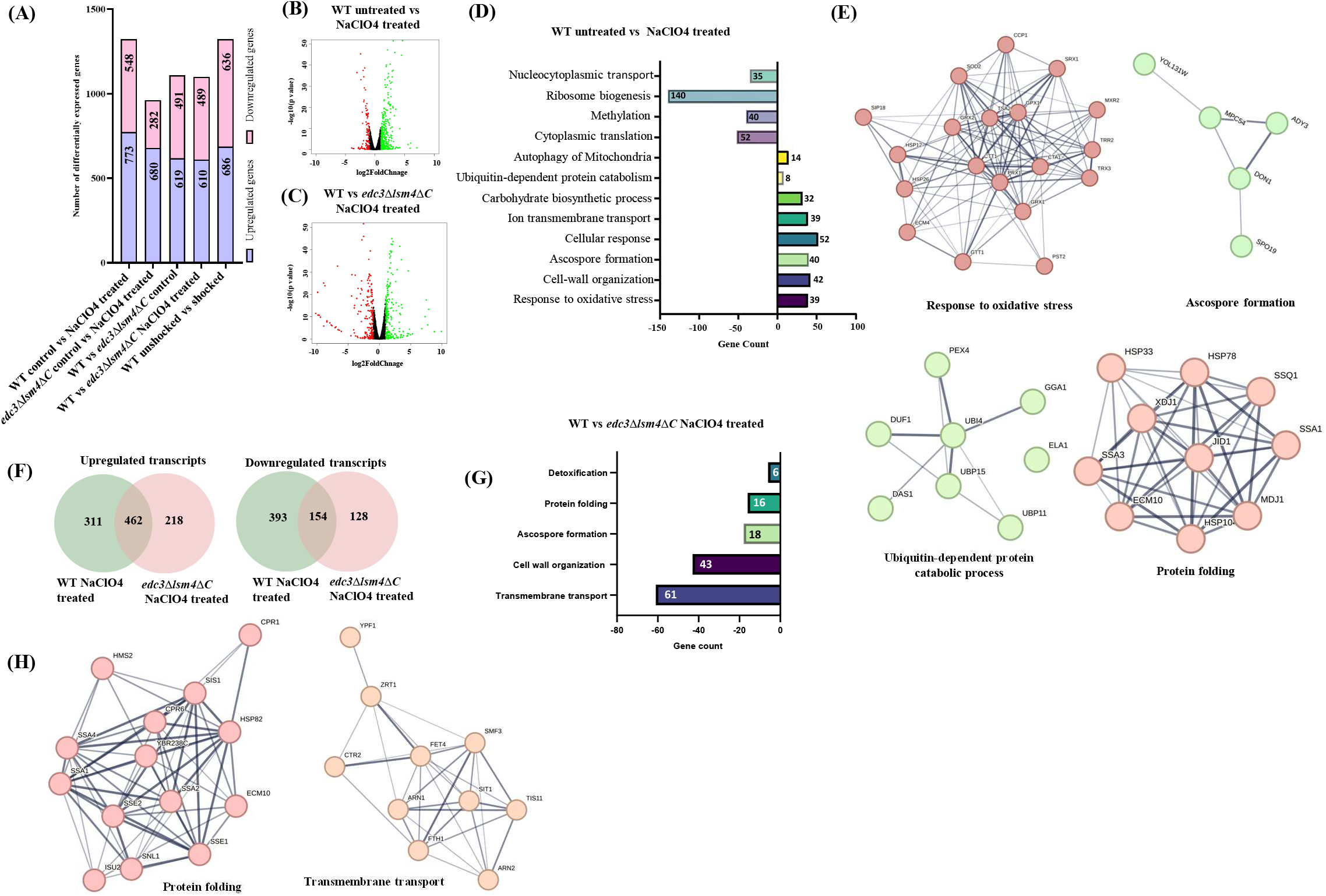
Transcriptomic analysis in wildtype and edc3Δlsm4ΔC reveals differential gene expression in response to sodium perchlorate. [A] Graph depicting the summary of differentially expressed genes in different conditions. [B-C] Volcano plot showing changes in transcripts level in [B] wildtype cells upon sodium perchlorate treatment, [C] wildtype cells upon edc3Δlsm4ΔC mutant under sodium perchlorate treatment. The green dots represent up-regulated transcripts, the red dots represent down-regulated transcripts and neutrally expressed transcripts are represented as black. [D] Gene enrichment analysis based on the biological process of differentially expressed genes in wild-type cells upon sodium perchlorate treatment. The X-axis represents the number of genes; a positive x-axis refers to upregulated transcripts, and a negative x-axis represents downregulated transcripts. [E] String analysis for 4 upregulated biological processes enriched in wild-type cells upon sodium perchlorate treatment. [F] Venn diagrams showing several differentially expressed transcripts upon sodium perchlorate treatment in wild-type and edc3Δlsm4ΔC cells. [G] Gene enrichment analysis based on the biological process of downregulated transcripts in edc3Δlsm4ΔC cells upon sodium perchlorate treatment compared to wild-type cells upon sodium perchlorate treatment. A negative x-axis represents downregulated transcripts [H] String analysis for a downregulated biological process enriched in edc3Δlsm4ΔC cells upon sodium perchlorate treatment.

RNA sequencing (RNA-seq) analysis of wild-type yeast cells treated with 100 mM sodium perchlorate identified 1,321 differentially expressed genes (DEGs), of which 773 were upregulated and 548 were significantly downregulated. Gene Ontology (GO) enrichment analysis of the upregulated genes revealed significant enrichment in several biological processes. Notably, genes associated with cellular respiration represented the largest category of upregulated genes in sodium perchlorate-treated wild-type cells (Figure 4D-E). Other enriched categories among the upregulated genes included those involved in oxidative stress response, cell wall organization, carbohydrate metabolism, ion transport, and mitochondrial autophagy. Additionally, genes involved in ascospore formation and ubiquitin-dependent protein degradation processes were also significantly upregulated. Conversely, the downregulated genes were enriched in processes related to cytoplasmic translation, methylation, nucleocytoplasmic transport, and ribosome biogenesis (Figure 4D). These findings suggest that sodium perchlorate might induces a metabolic shift, promoting stress adaptation while suppressing translation-related processes that are energy-intensive.

In the *edc3Δlsm4ΔC* mutant, sodium perchlorate treatment altered gene expression, with some differentially expressed genes overlapping with those in perchlorate-treated wild-type cells. 462 upregulated genes and 154 downregulated genes were common between wild-type and mutant cells treated with sodium perchlorate, highlighting a perchlorate-specific stress response (Figure 4F).

Comparison of the *edc3Δlsm4ΔC* mutant to wild-type under sodium perchlorate stress revealed 1,099 DEGs, with 610 genes upregulated and 489 genes significantly downregulated. GO enrichment analysis of the downregulated genes in the mutant highlighted significant enrichment of mRNAs encoding proteins involved in processes such as cell wall organization, protein folding, detoxification, transmembrane transport, and ascospore formation (Figure 4 G-H). This suggests that the *edc3Δlsm4ΔC* mutant likely exhibits an impaired response, particularly in pathways critical for maintaining cellular integrity and stress adaptation under perchlorate stress. This is consistent with the observed growth defects in the *edc3Δlsm4ΔC* mutant under continuous sodium perchlorate stress (Figure 3C-D).

To further elucidate the alteration of the global transcriptome during recovery from the shock wave, transcriptomic analysis was performed on wild-type *S. cerevisiae* recovering from exposure to 5.6 M intensity shock waves. Direct analysis of shocked cells was not possible due to technical limitations of the shock tube, which yielded only a small sample amount. Since shocked cells demonstrated a marked lag phase, we were curious to identify the transcriptome changes just before the cells moved out of the extended lag phase induced by shock waves. Transcriptome analysis revealed differential expression of 1,322 genes. Among these, 686 genes were upregulated, and 636 genes were downregulated (Figure 4A). Gene Ontology (GO) enrichment analysis of the upregulated genes highlighted significant enrichment in processes such as methylation, translation, and nuclear export (Supplementary Figure 5). Conversely, the downregulated genes were enriched in pathways associated with glucose import and cellular respiration. Notably, genes involved in oxidative stress response and temperature stimulus were also downregulated, which is intriguing as these pathways are commonly associated with stress adaptation (Supplementary Figure 5). This suggests that the recovery from shock waves might involve a distinct regulatory mechanism prioritizing specific cellular processes over generalized stress responses. The *edc3Δlsm4ΔC* mutant was not subjected to transcriptome analysis upon shock wave exposure, as we did not observe a significant difference in growth rate compared to wild-type shocked cells (Figure 3C-D)

**Figure 5:**
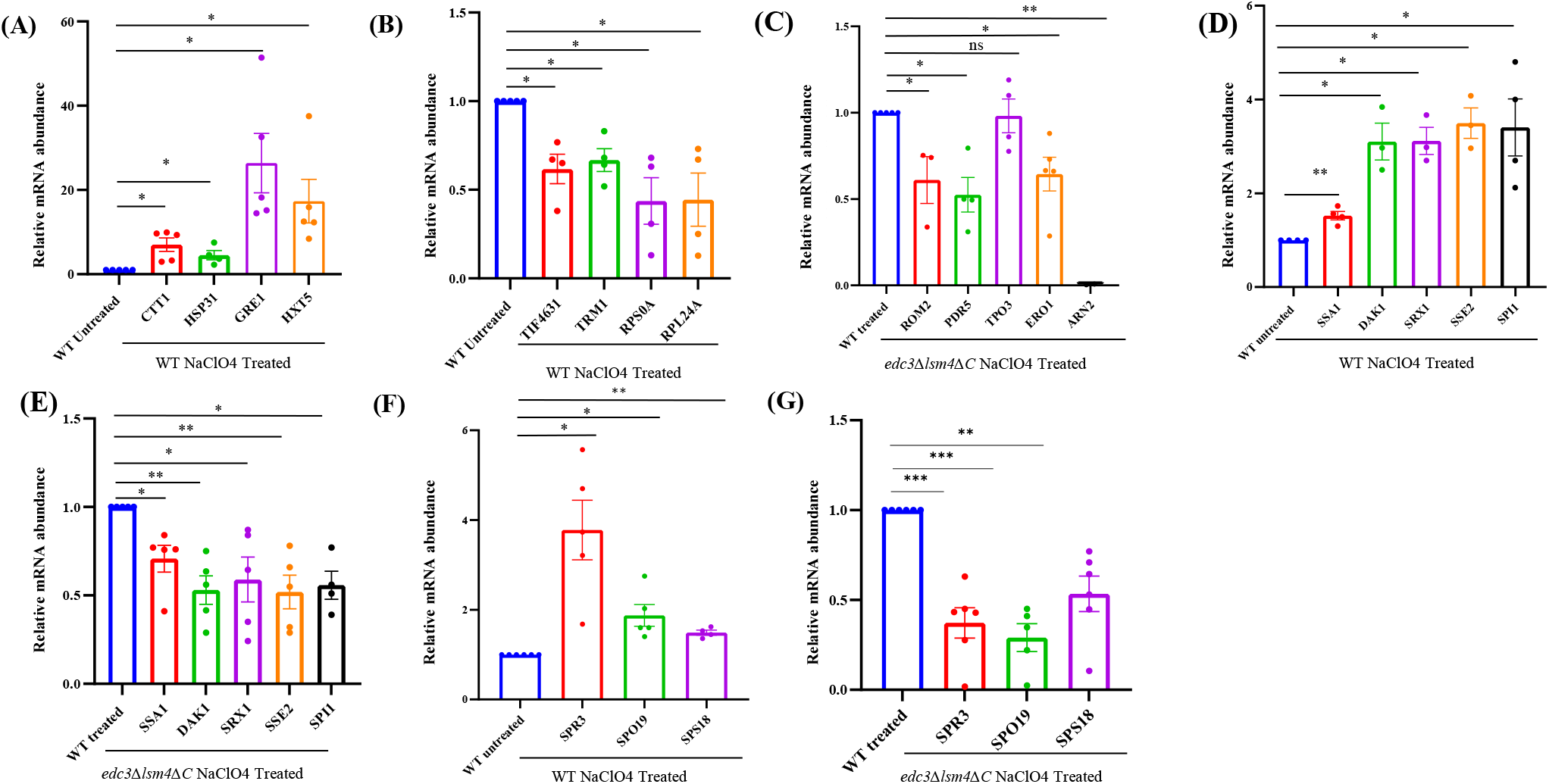
Validation of differentially expressed genes by qRT-PCR analysis. Validation of genes by qRT-PCR [A] upregulated in wildtype untreated vs sodium perchlorate treated; [B] downregulated in wildtype untreated vs sodium perchlorate treated; [C] neutrally expressed in wildtype treated but downregulated in *edc3Δlsm4ΔC* sodium perchlorate treated condition. [D-E] Validation of genes upregulated in wildtype treated [D] but downregulated in *edc3Δlsm4ΔC* sodium perchlorate treated condition [E] by qRT-PCR. [F-G] Validation of ascospore-related genes upregulated in wildtype treated [F] but downregulated in edc3Δlsm4ΔC sodium perchlorate treated condition [G] by qRT-PCR. Each data point represents biological replicate. For all validations, PGK1 was used as internal control. In the above experiments, error bars represent the Standard Error of the Mean. The significance of the data was calculated by paired t-test and p-values are summarized as: ^***^, p < 0.005; ^**^, p < 0.01; ^*^, p < 0.05 *. log2FC values obtained from RNA-seq for all genes validated are shown in Supplementary Figure 6.

### Validation of the transcriptome data

To select an appropriate internal control for validating the RNA-seq results, we assessed the mRNA levels of 3 reference genes-PGK1, ACT1, and SCR1 using qRT-PCR under the same conditions applied in the RNA-seq experiments. Among these, PGK1 and ACT1 exhibited no change in expression across all tested conditions, while SCR1 showed variable expression upon perchlorate treatment (Supplementary Figure 6A). Therefore, PGK1 was chosen as the internal control for mRNA target validation. To validate the differentially expressed transcripts identified through RNA-seq in wild-type yeast upon sodium perchlorate treatment, we focused on *CTT1, GRE1, HXT5*, and *HSP31*, which were upregulated, and *TIF4631, TRM1, RPS0A*, and *RPL24A*, which were downregulated. These transcripts were selected as representatives from each enriched GO category for qRT-PCR analysis. Notably, *GRE1* was among the transcripts that exhibited the highest fold change in RNAseq analysis. The qRT-PCR results validated the RNA-seq findings, with all tested transcripts exhibiting trends consistent with the sequencing data. The upregulated genes, including *CTT1* and *GRE1*, which are associated with oxidative stress response, and *HXT5*, linked to carbohydrate metabolism, showed increased mRNA levels. Similarly, *HSP31*, which is involved in stress protection, was also significantly upregulated (Figure 5A). Conversely, genes, such as *TIF4631* and *TRM1*, associated with translation initiation and methylation, respectively, exhibited reduced mRNA levels. Ribosomal protein genes *RPS0A* and *RPL24A* also showed decreased expression, aligning with the observed downregulation of translation-related processes (Figure 5B).

**Figure 6:**
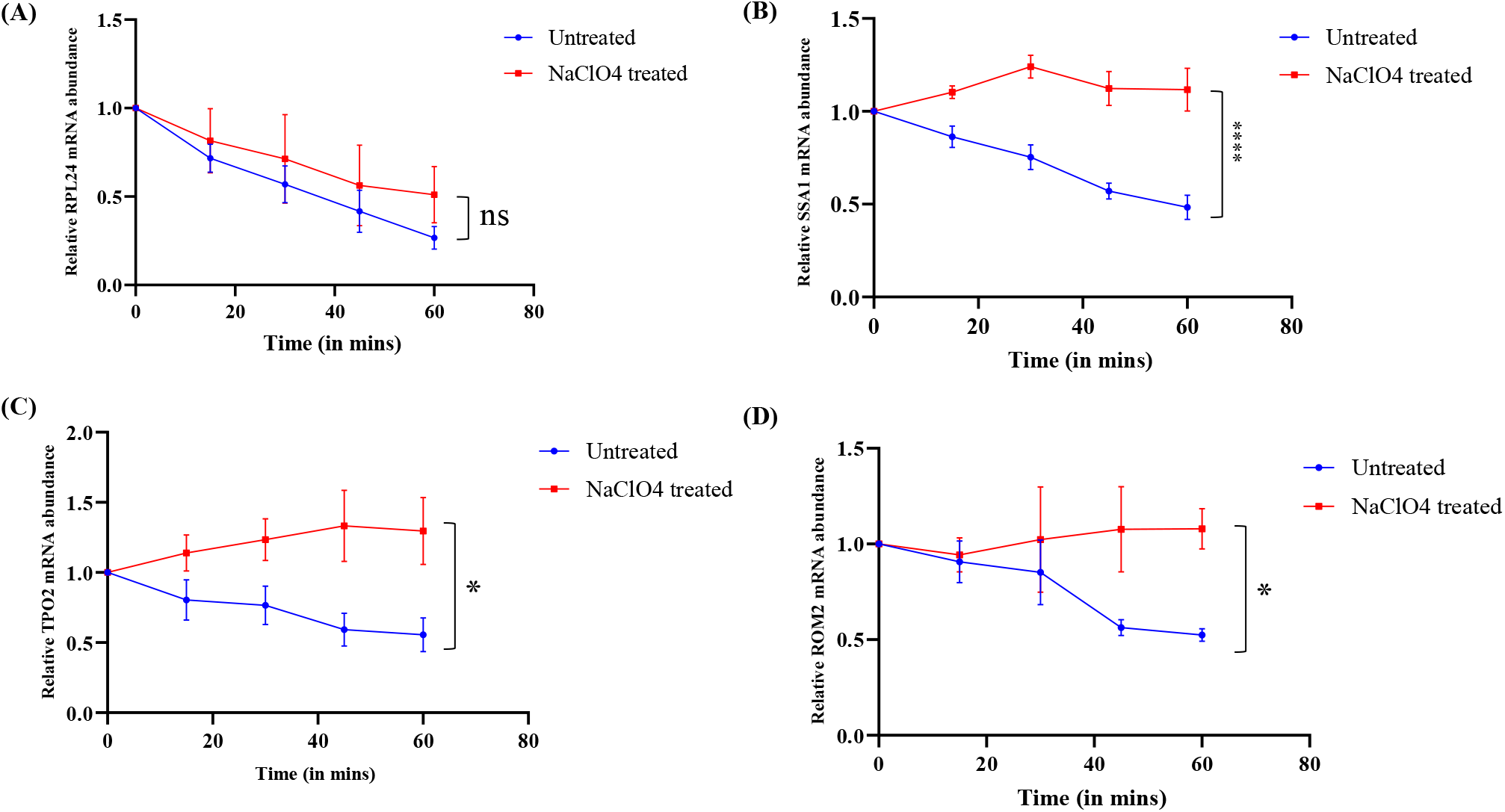
Stabilization of transcripts in wildtype upon sodium perchlorate treatment. mRNA stability assay by qRT-PCR in *rpb1-1* untreated and sodium perchlorate treated conditions for [A] RPL24; [B] SSA1; [C] TPO2; [D] ROM2 (n=3 biological replicates). For all validations PGK1 was used as internal control. In the above experiments, error bars represent Standard Error of the Mean. The significance of the data was calculated by 2-way ANOVA using Šídák’s multiple comparisons test and *p-*values are summarized as: ^****^, *p <* 0.0001; ^*^, *p <* 0.05.

Among the transcripts downregulated in *edc3Δlsm4ΔC* perchlorate-treated condition compared to wild-type perchlorate-treated, we validated the six transcripts. *ROM2, SSA1, PDR5, TPO3, ERO1, ARN2* transcripts (Figure 5C). These transcripts do not change in response to perchlorate in wild-type compared to untreated, but are significantly downregulated in *edc3Δlsm4ΔC* perchlorate-treated condition compared to wild-type perchlorate-treated, suggesting that these may be destabilized in the absence of RNP condensates. We also validated four transcripts, such as *DAK1, SRX1, SSE2,* and *SPI1,* which were upregulated in wild-type treated compared to untreated, but significantly downregulated in *edc3Δlsm4ΔC* perchlorate treated condition. Of all tested transcripts, except *TPO3*, all transcripts were validated (Figure 5-D-E). Furthermore, we validated three transcripts involved in ascospore wall assembly-*SPR3, SPO19,* and *SPS18* – which were upregulated in wildtype upon sodium perchlorate treatment but are significantly downregulated in *edc3Δlsm4ΔC* mutant under perchlorate stress (Figure 5F-G).

Similarly, to validate RNA sequencing result for differentially expressed mRNAs in wild-type during recovery from exposure to 5.6 M intensity shock waves, four mRNAs such as *TIF4631, NSR1, TRM1, HMT1* were chosen as representatives from upregulated GO transcripts. *TIF4631* was selected as it encodes a translation initiation factor crucial for translation regulation, while *HMT1* was chosen as it is a major arginine methyltransferase. The following five transcripts *CTT1, HXT5, SCO1, SDH1,* and *COX4* were chosen to validate downregulated mRNAs (Supplementary Figure 5E-F). *CTT1* and *HXT5* were among the transcripts that exhibited the lowest expression levels in RNA seq analysis. We observed that all tested transcripts except *COX4* validated the changes observed in RNA-seq analysis. Taken together, these observations validate the RNA-sequencing results by qRT-PCR and highlight the involvement of specific pathways that may facilitate adaptation in response to Martian stress conditions.

### Stabilization of transcripts correlates with the induction of P-bodies upon sodium perchlorate treatment

We hypothesised that the reduced abundance of transcripts in the *edc3Δlsm4ΔC mutant* could be a result of post-transcriptional regulation due to the absence of P-bodies. To investigate this, we performed a transcription shut-off assay using a temperature-sensitive mutant of RNA polymerase II, *rpb1-1*. Upon heat shock, RNA polymerase II becomes inactive, halting mRNA synthesis, enabling us to asses mRNA stability (Nonet et al., 1987; Parker et al., 1991). We assessed the mRNA decay profile in untreated and NaClO4-treated cells over a 60-minute time course post-heat shock to assess how transcript stability is affected by sodium perchlorate. When *rpb1-1* untreated and sodium perchlorate-treated conditions were subjected to transcription shut-off by heat shock, we observed stabilization of specific transcripts such as *SSA1* (involved in protein folding), *TPO2* (involved in ion transport), *ROM2* (involved in cell wall signaling) in sodium perchlorate-treated condition compared to untreated condition (Figure 6B-D) whereas *RPL24* which was used as a negative control shows a steady decrease in abundance over time in both untreated and treated condition. These results support the notion of stabilization of stress-responsive transcripts and correlation with induction of P-bodies under sodium perchlorate treatment.

## Discussion

Our study focuses on the impact of Martian stress conditions, such as shock waves and perchlorate stress, on *Saccharomyces cerevisiae.* Our study makes the following conclusions: 1) wild-type yeast survives exposure to 5.6 Mach intensity shock waves with an extended lag phase (Figure 1B-C), 2) wild-type yeast grows in presence of continuous 100mM sodium perchlorate with a significant growth defect compared to wildtype (Figure 1D-E), 3) Shock waves induces the assembly of RNP condensates (Figure 2A-C), 4) Sodium perchlorate induces P-body assembly (Figure 2D-E), 5) *edc3Δlsm4ΔC* mutant fails to assemble RNP condensates in response to either shock waves or sodium perchlorate (Figure 3 A-B), 6) *edc3Δlsm4ΔC* mutant exhibits decreased and delayed growth compared to wild-type in response to sodium perchlorate stress alone and shock waves combined with perchlorate stress (Figure 3C-D and Supplementary Figure 2 respectively), 7) Transcriptome data indicates that sodium perchlorate treatment triggers a metabolic shift in yeast, likely enhancing stress adaptation by ascospore formation while downregulating translation-related processes (Figure 4D), 8) Several transcripts are stabilized during perchlorate treatment, a condition under which P-body assembly is also induced in wildtype. (Figure 6) 9) *edc3Δlsm4ΔC* mutant exhibits an impaired stress response under sodium perchlorate stress, with downregulation of key transcripts associated with cellular integrity, protein folding and ascospore formation (Figure 4G-H), 10) Recovery from shock waves exposure triggers transcriptional changes in yeast including increased expression of genes related to translation and methylation, alongside reduced expression of transcripts associated with oxidative stress and metabolic pathways. (Supplementary Figure 5). These results identify an exciting possible connection between granule assembly and survival under Martian conditions.

Shock waves have been widely used for therapeutic applications (Rola et al., 2022). Use of shock waves for the destruction of kidney stones has been well-established (Chaussy et al., 1980). Shock wave therapy has also been used for various other therapeutic applications, such as in orthopedics for treating plantar fasciitis (Sun et al., 2017), in cardiology as an effective strategy for coronary artery disease (Nishida et al., 2004; Tepeköylü et al., 2017; Vainer et al., 2016), in neurology to reduce lower limb spasticity in stroke patients (Cabanas-Valdés et al., 2020) and for peripheral nerve regeneration (Sağir et al., 2019). Interestingly, beyond therapeutic uses, recent studies reveal the synthesis of amino acids from simple molecules in response to shock processes, which mimics the impact-induced shock events, highlighting the potential role of shock waves in the origin of life forms (Singh et al., 2020). However, the strength of shock waves used for therapeutic purposes in these studies is much weaker, around 0.09 mJ/mm^2^, corresponding to 2 bars. We hypothesized that shock waves of higher intensities could be stressful for lifeforms on Earth, as certain reports have shown shock waves to be detrimental (Li et al., 2014; Ohmori et al., 1994). To mimic the Martian conditions, we tested the ability of yeast cells to survive higher Mach value shockwaves generated by shock tubes in laboratory conditions. In our experiments, yeast survived and grew after exposure to 5.6M shock waves (Figure 1B), which was surprising. To the best of our knowledge, the survival of any model systems to such high-intensity shock waves has not been reported. Similar and much higher intensity shockwaves could be generated on Mars surface due to the impact of micrometeorites (Kurosawa et al., 2018). Our study highlights the credentials of yeast as a model system for studying survival under harsh Martian conditions in the future. However, upon exposure to the increasing intensity of shock waves, there was an increase in the lag phase, indicating that shock waves affect the cell physiology and induce a stress response. The ability to survive such high-intensity shock waves also highlights the presence of a robust repair mechanism and stress response pathway in yeast, making it capable of mitigating the impacts of sudden pressure and temperature changes of such high magnitude.

Interestingly, exposure to 5.6 Mach intensity shock waves in wild-type yeast induced the formation of P-bodies and stress granules (Figure 2A-C). A recent study focused on understanding the impact of specific simulated microgravity conditions on yeast. A major group of genes belonging to RNP condensate-related proteins was among the subset of genes that were reported to be differentially expressed in response to microgravity (Nemoto et al., 2019). This aligns with our observation, highlighting the involvement of RNP condensate in response to such stresses.

Our findings indicate that wild-type yeast can survive perchlorate at a concentration equivalent to that found on Martian soil with a slower growth rate than in unstressed conditions (Figure 1D). This growth defect could possibly be due to the chaotropic properties of perchlorate on biomolecules and their metabolic processes. In addition to its impact on growth, we observe that 100 mM sodium perchlorate induced the formation of P-bodies in wild-type yeast (Figure 2D-E). Unlike shock waves, perchlorate stress did not induce the formation of stress granules. This differential response indicates that the type of RNP condensates formed is specific to the nature of the Martian stress. These RNP condensates are known to play important roles in cellular stress responses by regulating mRNA fate at the level of mRNA stability and translation to alter the proteome (Liboy-Lugo et al., 2024; Marcelo et al., 2021). The formation of condensates suggests that yeast cells activate a conserved stress response pathway to cope with the damage caused by sodium perchlorate.

The *edc3Δlsm4ΔC* mutant, which is defective in assembly of RNP condensates, exhibited delayed growth compared to the wild-type in response to sodium perchlorate (Figures 3C-D). This delay in growth likely highlights the importance of RNP condensates in adaptation to stressful conditions like perchlorate. Considering that Martian conditions involve a combination of stresses, the combined effect of 5.6 Mach intensity shock waves and 100mM sodium perchlorate on the growth of wild-type and *edc3Δlsm4ΔC* mutant strains was also assessed. The wild-type strain exhibited slower growth under combined stress compared to individual stresses, and this growth retardation was more pronounced in the *edc3Δlsm4ΔC* mutant (Supplementary Figure 2). The combined stress of shock waves and sodium perchlorate worsens the growth defect in the mutant, indicating that multiple simultaneous stresses present a more significant challenge to cellular survival and adaptation. Assessing the impact of other Martian stress individually and in combination would be an important future direction. Interestingly, *edc3Δlsm4ΔC* mutant also demonstrates decreased survival post-continuous sodium perchlorate treatment (Figure 3G-H). This observation further highlights the possible critical role of RNP condensates in facilitating recovery from perchlorate stress. In contrast to perchlorate stress and combined stress, exposure to shock waves alone did not cause any growth defect for *edc3Δlsm4ΔC* mutant, indicating the specific role of condensates in regulating specific stress responses.

Our study provides some insights into the molecular mechanisms underlying yeast adaptation to Martian-like stress conditions, specifically sodium perchlorate and shock wave stress. Combining RNA sequencing and qRT-PCR validation, we identified key transcriptome changes and pathways likely involved in the cellular response to these stressors. RNA-seq analysis revealed that wild-type *S. cerevisiae* exposed to 100 mM sodium perchlorate undergoes extensive transcriptome remodelling, with changes in the abundance of 1,321 transcripts. Upregulated transcripts were enriched inprocesses such as cellular respiration, oxidative stress response, carbohydrate metabolism, and mitochondrial autophagy, indicating a robust adaptive response. These findings suggest that wild-type cells engage in metabolic and stress-response pathways to mitigate the detrimental effects of sodium perchlorate. For instance, the upregulation of 39 transcripts involved in oxidative stress response, including *CTT1* and *GRE1*, validated by qRT-PCR, may indicate activation of oxidative stress defence mechanisms. However, further work is needed to determine the extent to which these changes correspond to functional outcomes. Similarly, increased expression of *HXT5*, a gene linked to carbohydrate metabolism, may reflect enhanced glucose utilization.. We also observed the upregulation of cell wall organization-related genes in mitochondrial autophagy ria suggesting that perchlorate stress may impact cell wall and mitochondria. The upregulation of ubiquitin-dependent protein degradation process suggests that perchlorate stress induces protein damage or misfolding, requiring enhanced proteasomal activity to clear toxic protein aggregates or misfolded proteins and maintain cellular integrity to survive under perchlorate stress. This is consistent with the proteomic level changes in response to perchlorate reported in a halotolerant yeast *D. hansenii* (Heinz et al., 2022). The upregulation of genes *SPR3, SPO19, SPS18* involved in ascospore formation is intriguing. Since ascospore formation allows yeast cells to enter a stress-resistant dormant state, it is possible that *S. cerevisiae* initiates spore formation to adapt to perchlorate stress. Sporulation is typically induced under nutrient deprivation (Honigberg & Purnapatre, 2003; Neiman, 2011), this further highlights that perchlorate stress might trigger a broader cellular response mimicking nutrient deprivation, leading to the activation of the sporulation process. Notably, genes such as NDT80, a key regulator for the sporulation process known to induce downstream transcriptional targets involved in the assembly of prospore membrane and spore wall (Chu et al., 1998; Winter, 2012), are also upregulated in wild-type upon perchlorate treatment. The upregulation of *NDT80* suggests that the yeast cells may be attempting to undergo a developmental transition to enhance survival under adverse conditions. The laboratory strains, including BY4741, used here are haploid strains; however, our results argue that a diploid yeast might undergo sporulation in response to perchlorate stress.

Interestingly, the downregulated transcripts were enriched in processes such as cytoplasmic translation and ribosome biogenesis. This may indicate suppression of energy-intensive processes such as translation. Our observation is consistent with reports on other stresses in *S. cerevisiae*, which lead to global translation repression (Ashe et al., 2000; Garg et al., 2020; Melamed et al., 2008; Shenton et al., 2006). The reduced abundance of ribosomal protein genes *RPS0A* and *RPL24A*, as validated by qRT-PCR, supports this possibility, of survival over growth (which requires active translation and, therefore, energy) during stress. These observations indicate that the response to sodium perchlorate stress is similar to other oxidative stress response pathways.

In contrast, the *edc3Δlsm4ΔC* mutant, defective in RNP condensate assembly, exhibited a markedly impaired response to sodium perchlorate. Transcripts associated with key pathways, including cell wall organization, protein folding, and detoxification, were significantly downregulated in the mutant, likely suggesting an impaired ability to mount an effective stress response. These findings raise the possibility that RNP condensates, such as P-bodies, may contribute to transcript stabilization during stress, consistent with previous studies suggesting their role in mRNA storage and protection.(Hubstenberger et al., 2017; Standart & Weil, 2018). mRNA stability analysis (Figure 6) further support this notion, showing that transcripts like *SSA1, ROM2*, and *TPO2* are more stable in wild-type cells upon perchlorate exposure, a condition under which P-bodies are induced. This observation aligns with the hypothesis that in wild-type cells, RNP condensate formation may help buffer specific transcripts from degradation, thereby facilitating a more robust stress response.

However, it is important to note that both Edc3 and Lsm4 are RNA-binding proteins that could affect mRNA stability independently of P-bodies. Therefore, the *edc3Δlsm4ΔC* mutation might have pleiotropic effects beyond disrupting P-body assembly. This is a general caveat that applies to all studies using such mutants (Decker et al., 2007), and it is challenging to rescue the PB assembly defect in this background to assess the pleiotropic effects resulting from this mutant background. Therefore, while our data are consistent with a role for P-bodies in transcript stabilization and stress resilience, we cannot rule out additional effects of the *edc3Δ lsm4ΔC* mutation on cell physiology, mRNA decay pathways, or stress response regulation.

Having said that, this study provides a robust framework to initiate functional analysis of condensates in response to Martian stress.

In wild-type yeast exposed to 5.6 Mach intensity shock waves, we observed a distinct mRNA abundance profile. The upregulation of genes involved in methylation, translation, and nuclear export, suggests likely activation of transcriptional and translational machinery post-shock wave exposure. This observation can be explained by the timing of RNA sequencing, as samples were collected during the recovery phase, capturing the cellular response during recovery rather than the immediate impact of shock waves. This likely reflects a cellular effort to restore homeostasis and initiate repair mechanisms. For example, the increased expression of *TIF4631* and *NSR1*, validated by qRT-PCR, may reflect a cellular attempt to restore translation and ribosome function, processes that are important for recovery under stress in general. Conversely, the downregulation of genes associated with glucose import (*HXT5*) and cellular respiration (*SDH1* and *SCO1*) might suggest a strategic conservation of energy during recovery. This downregulation may reflect a shift toward suppressing metabolic activity to prioritize repair processes. Intriguingly, genes typically associated with generalized stress responses, such as oxidative stress and temperature stimulus, were also downregulated during recovery. This likely suggests that the cellular response to shock waves involves a unique regulatory mechanism that diverges from canonical stress pathways. Nonetheless, the functional relevance of these transcript abundance changes requires further exploration.

The mRNA abundance profiles observed in response to sodium perchlorate and shock wave stress highlight the ability of yeast to adapt to diverse environmental challenges. The ability of wild-type cells to modulate energy metabolism, stress response pathways, and translational activity underscores the robustness of their adaptive mechanisms. In contrast, the impaired response in *edc3Δlsm4ΔC* mutant likely emphasizes the critical role of RNP condensates in coordinating these processes (Figure 7). These findings have broader implications for understanding the resilience of yeast under Martian conditions. Sodium perchlorate and shock waves represent just two of the many stressors present on Mars. The ability of yeast to survive and adapt to these stresses provides valuable insights into the potential for life to endure extraterrestrial environments. Moreover, the conserved nature of stress response pathways from yeast to humans suggests that these findings could inform strategies for utilizing RNP condensates as markers for assessing human health before and after space explorations, as stress-induced RNA granule assembly is a conserved feature of human cells.

**Figure 7:**
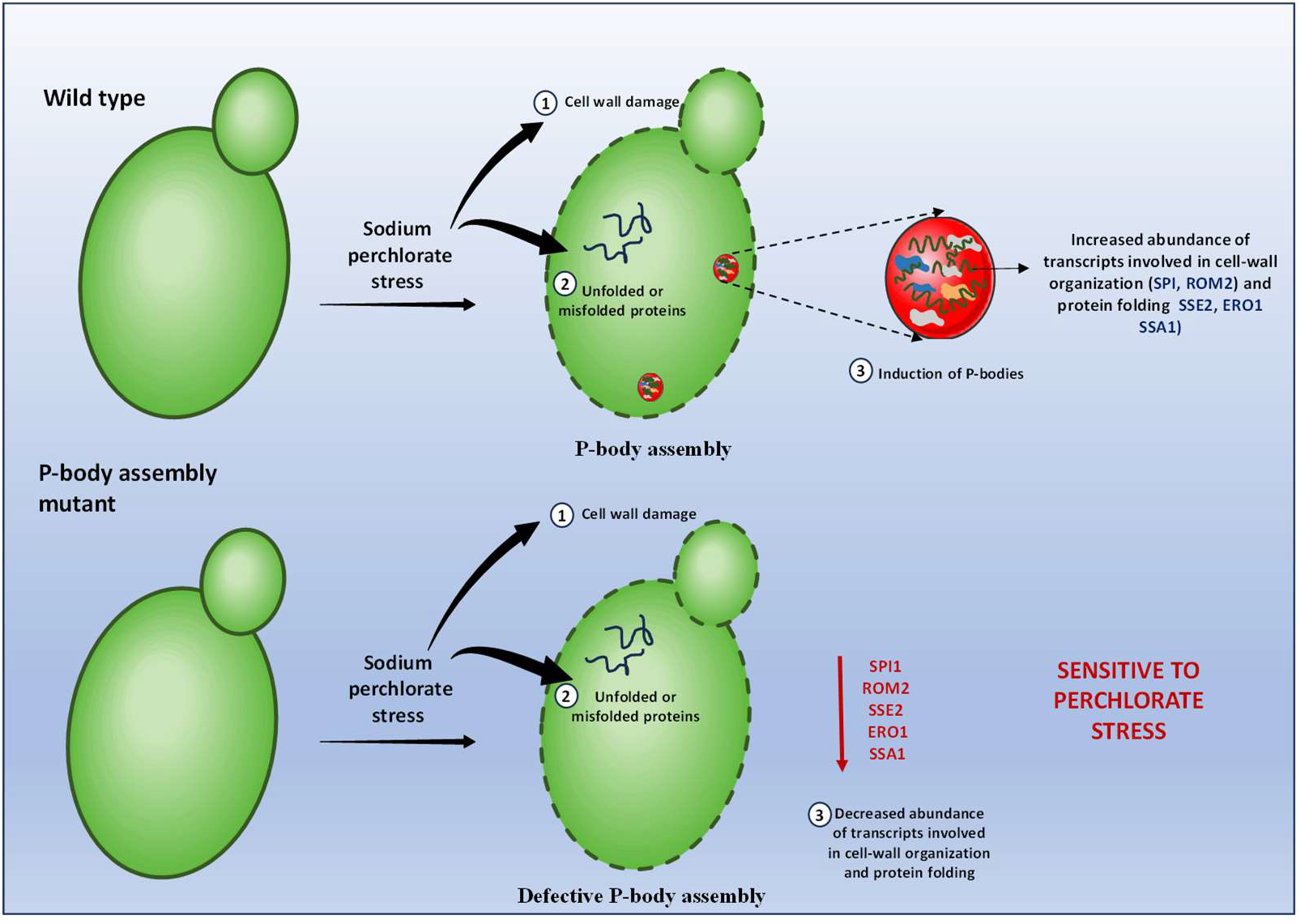
Model depicting the importance of RNP condensate in mediating survival under Mars-like stress condition.

## Supporting information

all supplementary tables

all supplementary figures

## Data Availability

The data discussed in this study has been deposited in NCBI’s Gene Expression Omnibus, and are accessible through GEO accession number: GSE293434. (https://www.ncbi.nlm.nih.gov/geo/query/acc.cgi?&acc=GSE293434)

## Acknowledgment

PIR and BS thank IISc-ISRO space technology cell (STC) STC/BES/PR/479 for the funding. RD thanks STC/BES/PR/479 for the project assistantship. The authors also thank CRG/2022/000594 for funding. The authors thank the Physical Research Laboratory (PRL) Department of Space, Government of India, for funding. Funding from DST-FIST is graciously acknowledged. We acknowledge Genotypic Technology Private Limited, Bangalore for NGS sequencing and data analysis reported in this study. We thank Vijay Thiruvenkatam, Dhiraj Bhatia and Sivapriya Kirubakaran at the Indian Institute of Technology, Gandhinagar, for providing logistical and infrastructure support to perform some of the experiments.

## Declaration of interests

The authors declare no competing interests.

## Notes

https://www.ncbi.nlm.nih.gov/geo/query/acc.cgi?&acc=GSE293434

### Competing Interest Statement

The authors have declared no competing interest.

