## Supplementary material for "Ribonucleoprotein (RNP) condensates modulate survival in response to Mars-like stress conditions": all supplementary tables

**Supplementary Table 1: Yeast strains used in this study.**

| Description | Genotype |
|---|---|
| Wild-type (BY4741) yeast cells | *MAT*a *his3D1 leu2 met15 ura3* (BY4741) |
| yPIR58 | MATα leu2 trp1 ura3 lys2 his4 cup1::LEU2/PGK1pG/MFA2pG Lsm4Dc::NEO edc3::NEO (*edc3*Δ*lsm4*Δ*C)* |
| yPIR141 | MATα ura3-52 his4-539 *rpb1-1* |

**Supplementary Table 2: Plasmids used in this study.**

| Plasmid Name | Description | Vector |
|---|---|---|
| pPIR50 | Used to express Dcp2-mCherry and Pab1 GFP protein under the own promoter (CEN) | pRS416 |
| pPIR51 | Used to express Scd6-GFP protein under the Scd6 promoter (CEN) | pRS316 |

**Supplementary Table 3: Primers used for qRT-PCR.**

| Gene | Sequence |
|---|---|
| CTT1 F | GTACACCAGACACTGCAAGA |
| CTT1 R | GAGGAAGAAGACGGGAGTATTG |
| HSP31 F | CCTTACATCCCTTCAACACATTC |
| HSP31 F | CCGTTTCGTCTTGACCATTTAG |

| | |
|---|---|
| GRE1 F | GTCAGTTGGACGACGATGAA |
| GRE1 R | GTCTACCGCTGCCGATATTAG |
| HXT5 F | GTTCGTGTTTGGGTGGGATA |
| HXT5 R | GGAAAGATAGGTAGTCCCGTTTG |
| ROM2 F | AATTCCGGGAACCACTCTTATC |
| ROM2 R | TGCTGGGACAATGAACTACTC |
| SSA1 F | CTCCATGGTCTTGGGTAAGATG |
| SSA1 R | TCCTTGGTAGCTTGTCTTTGAG |
| PDR5 F | ATTTCCGTGGTTCTGCTATGT |
| PDR5 R | CGTCAGCACTTGGATGGTATAA |
| TPO3 F | CTCGCTGGGTCCTTTGATTT |
| TPO3 R | GCTAAGGCGCAAGGGATATT |
| ERO1 F | GCAAATCCGGAACGATTTACAG |
| ERO1 R | GGCCAATGATTCACCAGTTTC |
| ARN2 F | GACGCAGTCATTGTACCCTAAA |
| ARN2 R | TTCAACCATCGCAGACCTAAC |
| DAK1 F | GGTGGTATGGTTGGTAGAAGAG |
| DAK1 R | CTGTACCGTCTAAGCCATACTTAC |
| SRX1 F | TGCTCTCTAGAACAGGCTGAG |
| SRX1 R | GCCGCCGAAGGCATAATATAG |
| SSE2 F | CTCTCCAACTTTAAGGGTCAGG |
| SSE2 R | TCCAAACGGTCTTCGTCATC |
| SPI1 F | TCGAGAAGCCTACTTCAGAAAC |
| SPI1 R | CATAACTGCACCAGCCAAAC |

| | |
|---|---|
| RRP45 F | GAAATGTTGTCACGAGGCTTAC |
| RRP45 R | CATGGCTGCATACTTGTTTCTC |
| NAB3 F | ACCTTCCACCTAACGTTGTATC |
| NAB3 R | CTGGCCTTGCATTGATTGTATT |
| TIF4631 F | CCAAGCTACAGTCTCAGGAAAG |
| TIF4631 R | GGGTAGGAGTTGGAGTAGAAGTA |
| NSR1 F | CCATCTGAACCATCTGACACTT |
| NSR1 R | ACTTCACCGTGTTTAGCGAATA |
| TIF3 F | AGAGGTGCTCAGTTTGGTAAG |
| TIF3 R | TGGTTGCTCATCAGTAGTCTTT |
| TRM1 F | GTATTGAGGAGGGTGGTCTAATG |
| TRM1 R | CGCACTTTCGTGAGTTGATTC |
| HMT1 F | GACGGCCAAGAGACAAGATATG |
| HMT1 R | CAGTGAGTGTATGGAGCATGAG |
| SCO1 F | GAACGCATGCAAGAAGTACAG |
| SCO1 R | CCTTCAGGGTCCATGAGATAAA |
| SDH1 F | GACGTCACCAAGGAACCTATTC |
| SDH1 R | CCTTGTCTTCGCCAGTTTCT |
| COX4 F | TTCAAGCCAGCCACAAGAA |
| COX4 R | TAGCACCAGGACCAATCAAAG |
| SPR3 F | GACAAAGTTCCCAAGGACAGTA |
| SPR3 R | GGTCCCTGTTCCTGCATTTA |
| SPO19 F | AACGACTACTATTCGGCCTTTC |
| SPO19 R | TCGCATTATCGTAGTCCTCATTC |

| | |
|---|---|
| SPS18 F | AGATTCGGTTCCTTCTTGTCC |
| SPS18 R | CCAGCCTCCTCTTGTAACTTT |
| AMA1 F | TTCTACAGAGGTTGACCGTTTC |
| AMA1 R | CGCTCAGGTGAAGGAGATTT |
| TPO2 F | CACTGTCCTGTTGTCCATCTT |
| TPO2 R | AAATAGCGGCTTCCATACCTAC |
| PGK1 F | ATGTCTTTATCTTCAAAGTTGT |
| PGK1 R | GGTTGGCAAAGCAGC |

**Supplementary Table 4: Statistical significance for all growth curve comparisions shown in Figure 1.**

| Figure | Comparisons | Statistical test used | Statistical significance | p-value |
|---|---|---|---|---|
| Figure 1B | WT (no shock) vs WT (5.6 M shock) | Tukey's multiple comparison test | **** | <0.0001 |
| Figure 1B | WT (no shock) vs WT (4.8 M shock) | Tukey's multiple comparison test | **** | <0.0001 |
| Figure 1B | WT (no shock) vs WT (3.5 M shock) | Tukey's multiple comparison test | ** | <0.005 |
| Figure 1B | WT (5.6 M shock) vs WT (4.8 M shock) | Tukey's multiple comparison test | * | <0.05 |
| Figure 1B | WT (5.6 M shock) vs WT (3.5 M shock) | Tukey's multiple comparison test | ** | <0.005 |
| Figure 1B | WT (4.8 M shock) vs WT (3.5 M shock) | Tukey's multiple comparison test | * | <0.05 |
| Figure 1C | WT (no shock) vs WT (5.6 M shock) | Tukey's multiple comparison test | **** | <0.0001 |
| Figure 1C | WT (no shock) vs WT (4.8 M shock) | Tukey's multiple comparison test | **** | <0.0001 |
| Figure 1C | WT (no shock) vs WT (3.5 M shock) | Tukey's multiple comparison test | * | <0.05 |
| Figure 1C | WT (5.6 M shock) vs WT (4.8 M shock) | Tukey's multiple comparison test | **** | <0.0001 |

| | | | | |
|---|---|---|---|---|
| Figure 1C | WT (5.6 M shock) vs WT (3.5 M shock) | Tukey's multiple comparison test | *** | <0.001 |
| Figure 1C | WT (4.8 M shock) vs WT (3.5 M shock) | Tukey's multiple comparison test | * | <0.05 |
| Figure 1D | Untreated vs. 50mM | Tukey's multiple comparison test | **** | <0.0001 |
| Figure 1D | Untreated vs. 100mM | Tukey's multiple comparison test | **** | <0.0001 |
| Figure 1D | Untreated vs. 200mM | Tukey's multiple comparison test | **** | <0.0001 |
| Figure 1D | Untreated vs. 500mM | Tukey's multiple comparison test | **** | <0.0001 |
| Figure 1D | Untreated vs. 1M | Tukey's multiple comparison test | **** | <0.0001 |
| Figure 1D | Untreated vs. 2M | Tukey's multiple comparison test | **** | <0.0001 |
| Figure 1D | 50mM vs. 100mM | Tukey's multiple comparison test | **** | <0.0001 |
| Figure 1D | 50mM vs. 200mM | Tukey's multiple comparison test | **** | <0.0001 |
| Figure 1D | 50mM vs. 500mM | Tukey's multiple comparison test | **** | <0.0001 |
| Figure 1D | 50mM vs. 1M | Tukey's multiple comparison test | **** | <0.0001 |
| Figure 1D | 50mM vs. 2M | Tukey's multiple comparison test | **** | <0.0001 |
| Figure 1D | 100mM vs. 200mM | Tukey's multiple comparison test | **** | <0.0001 |
| Figure 1D | 100mM vs. 500mM | Tukey's multiple comparison test | **** | <0.0001 |
| Figure 1D | 100mM vs. 1M | Tukey's multiple comparison test | **** | <0.0001 |
| Figure 1D | 100mM vs. 2M | Tukey's multiple comparison test | **** | <0.0001 |
| Figure 1D | 200mM vs. 500mM | Tukey's multiple comparison test | **** | <0.0001 |
| Figure 1D | 200mM vs. 1M | Tukey's multiple comparison test | **** | <0.0001 |

| | | | | |
|---|---|---|---|---|
| Figure 1D | 200mM vs. 2M | Tukey's multiple comparison test | **** | <0.0001 |
| Figure 1D | 500mM vs. 1M | Tukey's multiple comparison test | ns | 0.9553 |
| Figure 1D | 500mM vs. 2M | Tukey's multiple comparison test | ns | 0.1674 |
| Figure 1D | 1M vs. 2M | Tukey's multiple comparison test | ns | 0.7477 |
| Figure 1E | Untreated vs. 50mM | Tukey's multiple comparison test | ns | 0.8316 |
| Figure 1E | Untreated vs. 100mM | Tukey's multiple comparison test | ** | 0.0014 |
| Figure 1E | Untreated vs. 200mM | Tukey's multiple comparison test | **** | <0.0001 |
| Figure 1E | Untreated vs. 500mM | Tukey's multiple comparison test | **** | <0.0001 |
| Figure 1E | Untreated vs. 1M | Tukey's multiple comparison test | *** | 0.0004 |
| Figure 1E | 50mM vs. 100mM | Tukey's multiple comparison test | ** | 0.0039 |
| Figure 1E | 50mM vs. 200mM | Tukey's multiple comparison test | *** | 0.0002 |
| Figure 1E | 50mM vs. 500mM | Tukey's multiple comparison test | *** | 0.0001 |
| Figure 1E | 50mM vs. 1M | Tukey's multiple comparison test | *** | 0.0007 |
| Figure 1E | 100mM vs. 200mM | Tukey's multiple comparison test | * | 0.0350 |
| Figure 1E | 100mM vs. 500mM | Tukey's multiple comparison test | * | 0.0123 |
| Figure 1E | 100mM vs. 1M | Tukey's multiple comparison test | * | 0.0383 |
| Figure 1E | 200mM vs. 500mM | Tukey's multiple comparison test | ns | 0.7035 |
| Figure 1E | 200mM vs. 1M | Tukey's multiple comparison test | ns | 0.7949 |
| Figure 1E | 500mM vs. 1M | Tukey's multiple comparison test | ns | >0.9999 |
