## Supplementary material for "Ribonucleoprotein (RNP) condensates modulate survival in response to Mars-like stress conditions": all supplementary figures

### Slide 1
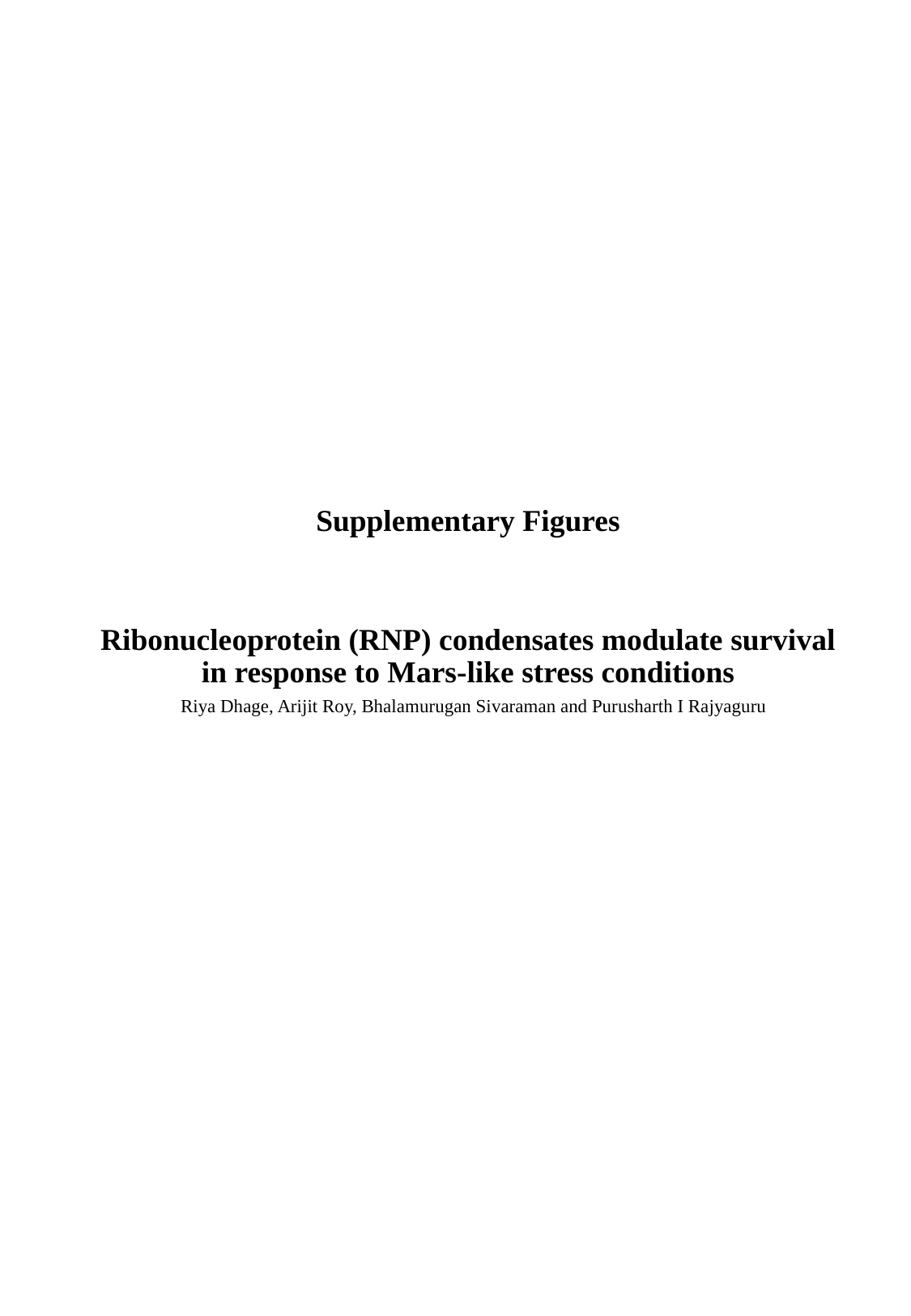

Ribonucleoprotein (RNP) condensates modulate survival in response to Mars-like stress conditions
Supplementary Figures
Riya Dhage, Arijit Roy, Bhalamurugan Sivaraman and Purusharth I Rajyaguru

### Slide 2
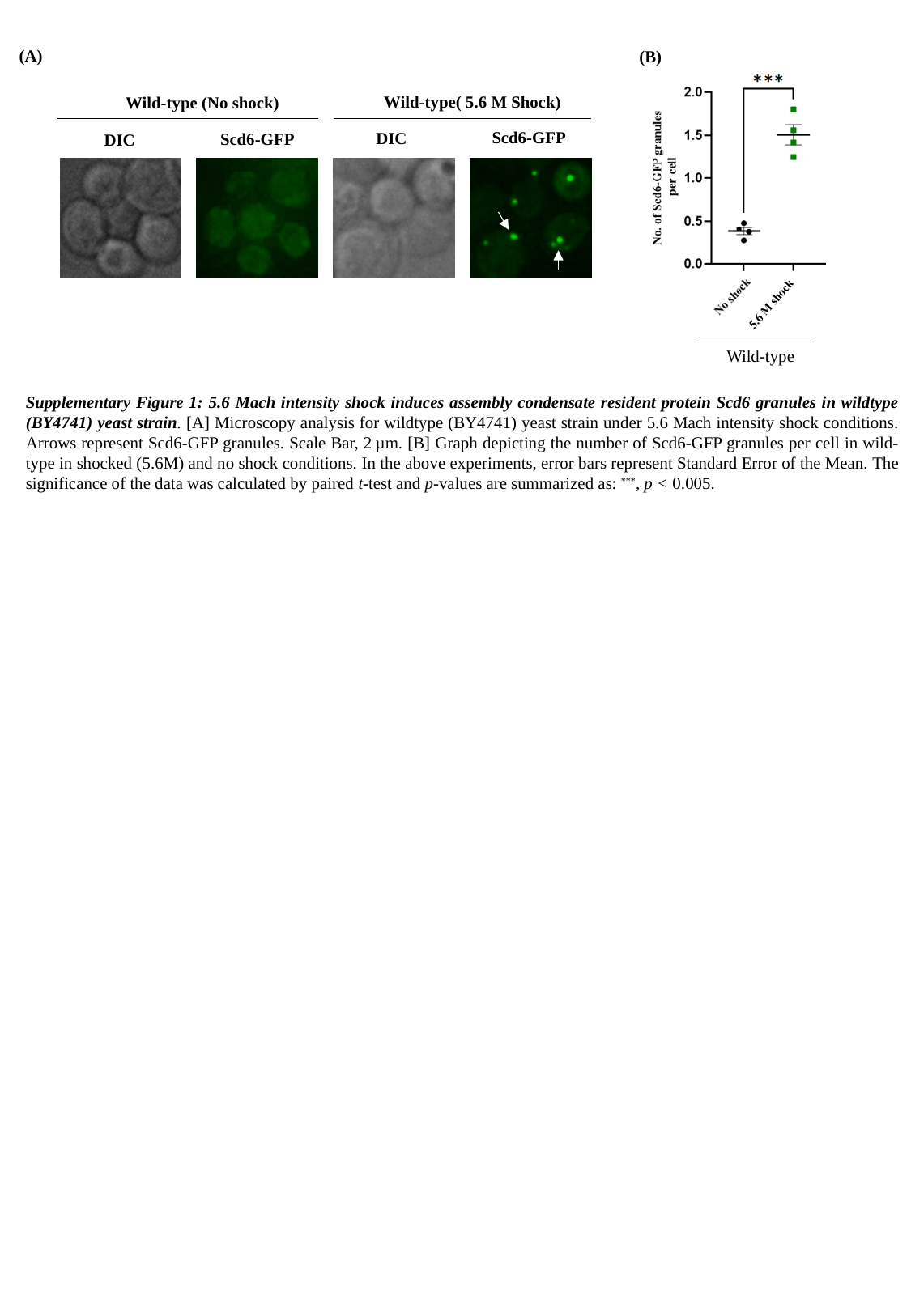

(A)
(B)
Wild-type( 5.6 M Shock)
Wild-type (No shock)
Scd6-GFP
DIC
Scd6-GFP
DIC
Wild-type
Supplementary Figure 1: 5.6 Mach intensity shock induces assembly condensate resident protein Scd6 granules in wildtype (BY4741) yeast strain. [A] Microscopy analysis for wildtype (BY4741) yeast strain under 5.6 Mach intensity shock conditions. Arrows represent Scd6-GFP granules. Scale Bar, 2 µm. [B] Graph depicting the number of Scd6-GFP granules per cell in wild-type in shocked (5.6M) and no shock conditions. In the above experiments, error bars represent Standard Error of the Mean. The significance of the data was calculated by paired t-test and p-values are summarized as: ^***^, p < 0.005.

### Slide 3
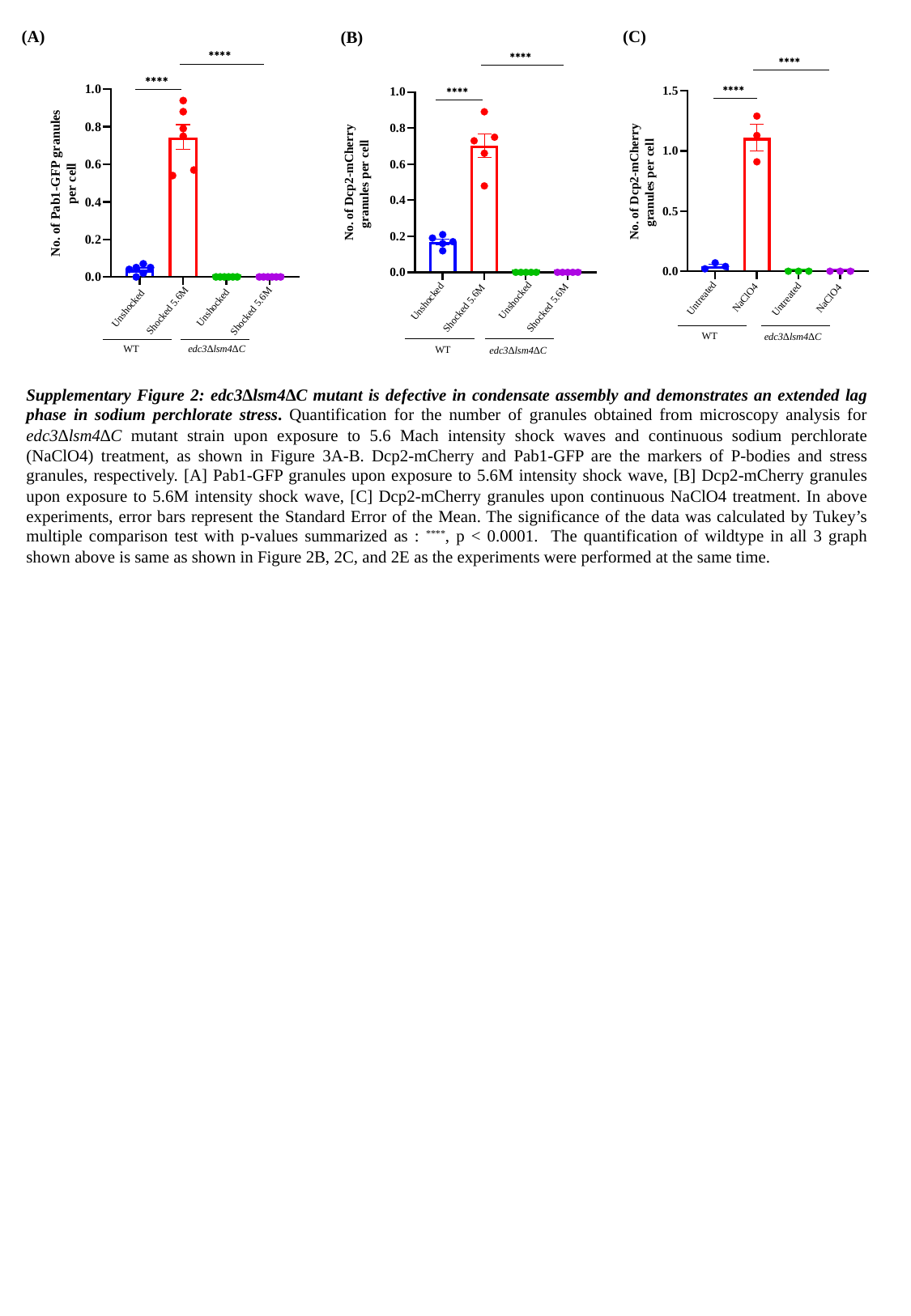

(C)
(A)
(B)
Unshocked
Unshocked
Shocked 5.6M
Shocked 5.6M
WT
edc3∆lsm4∆C
Unshocked
Unshocked
Shocked 5.6M
Shocked 5.6M
WT
edc3∆lsm4∆C
NaClO4
Untreated
Untreated
NaClO4
WT
edc3∆lsm4∆C
Supplementary Figure 2: edc3∆lsm4∆C mutant is defective in condensate assembly and demonstrates an extended lag phase in sodium perchlorate stress. Quantification for the number of granules obtained from microscopy analysis for edc3∆lsm4∆C mutant strain upon exposure to 5.6 Mach intensity shock waves and continuous sodium perchlorate (NaClO4) treatment, as shown in Figure 3A-B. Dcp2-mCherry and Pab1-GFP are the markers of P-bodies and stress granules, respectively. [A] Pab1-GFP granules upon exposure to 5.6M intensity shock wave, [B] Dcp2-mCherry granules upon exposure to 5.6M intensity shock wave, [C] Dcp2-mCherry granules upon continuous NaClO4 treatment. In above experiments, error bars represent the Standard Error of the Mean. The significance of the data was calculated by Tukey’s multiple comparison test with p-values summarized as : ^****^, p < 0.0001. The quantification of wildtype in all 3 graph shown above is same as shown in Figure 2B, 2C, and 2E as the experiments were performed at the same time.

### Slide 4
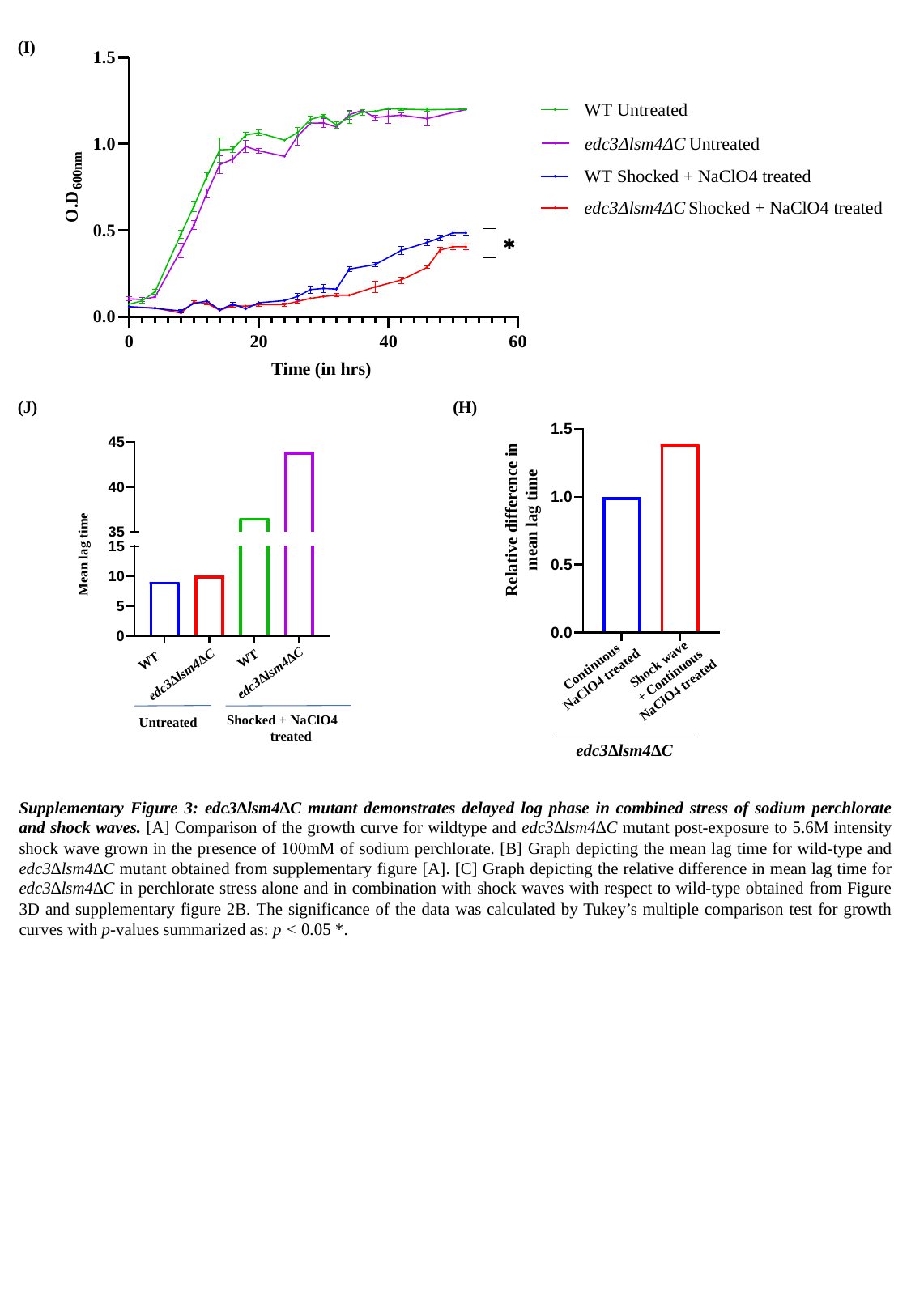

(I)
(H)
(J)
Relative difference in mean lag time
Shock wave
 + Continuous NaClO4 treated
Continuous NaClO4 treated
edc3∆lsm4∆C
Mean lag time
WT
WT
edc3∆lsm4∆C
edc3∆lsm4∆C
Shocked + NaClO4
 treated
Untreated
Supplementary Figure 3: edc3∆lsm4∆C mutant demonstrates delayed log phase in combined stress of sodium perchlorate and shock waves. [A] Comparison of the growth curve for wildtype and edc3∆lsm4∆C mutant post-exposure to 5.6M intensity shock wave grown in the presence of 100mM of sodium perchlorate. [B] Graph depicting the mean lag time for wild-type and edc3∆lsm4∆C mutant obtained from supplementary figure [A]. [C] Graph depicting the relative difference in mean lag time for edc3∆lsm4∆C in perchlorate stress alone and in combination with shock waves with respect to wild-type obtained from Figure 3D and supplementary figure 2B. The significance of the data was calculated by Tukey’s multiple comparison test for growth curves with p-values summarized as: p < 0.05 *.

### Slide 5
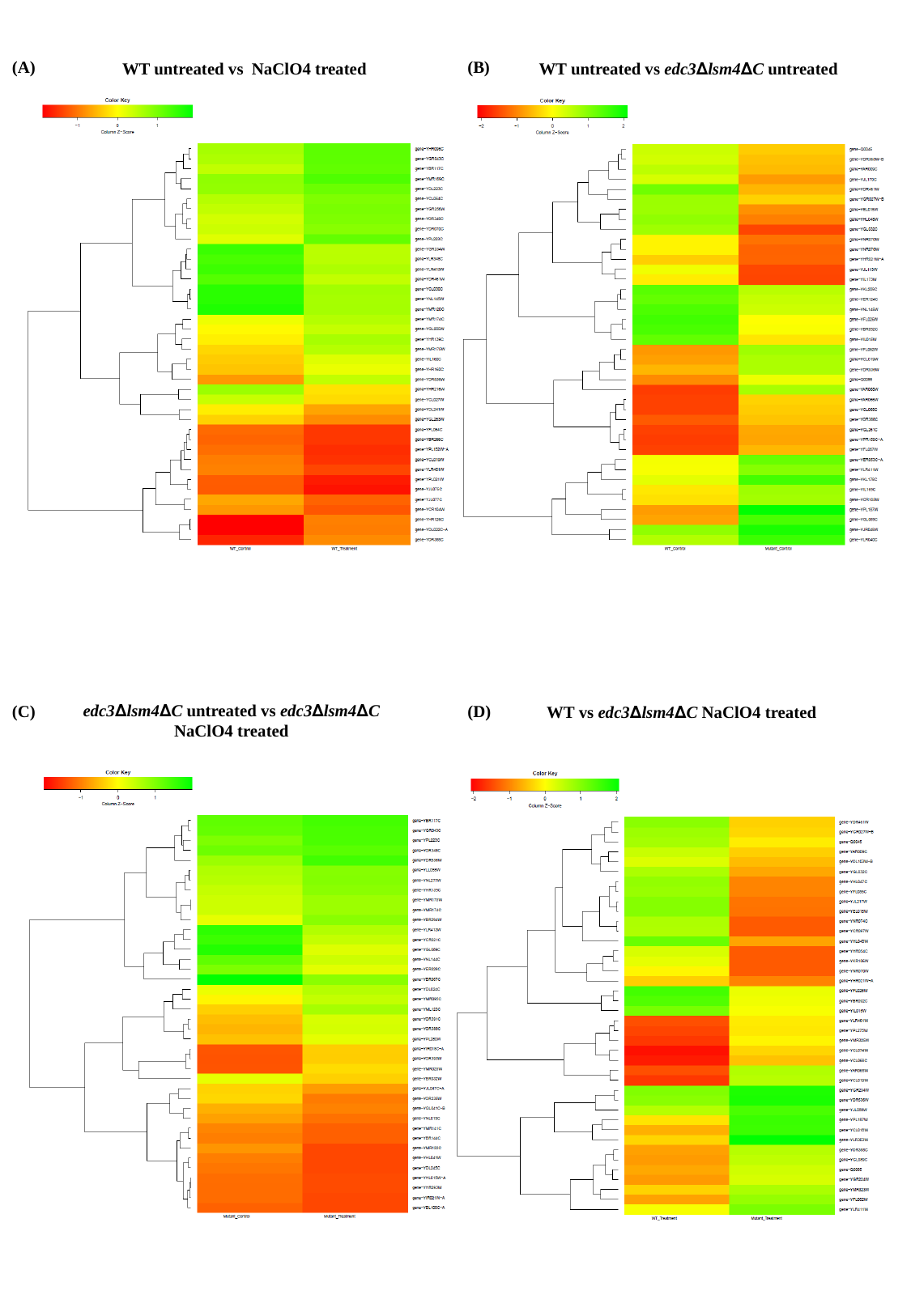

(B)
(A)
WT untreated vs NaClO4 treated
WT untreated vs edc3∆lsm4∆C untreated
 edc3∆lsm4∆C untreated vs edc3∆lsm4∆C
 NaClO4 treated
(D)
(C)
WT vs edc3∆lsm4∆C NaClO4 treated

### Slide 6
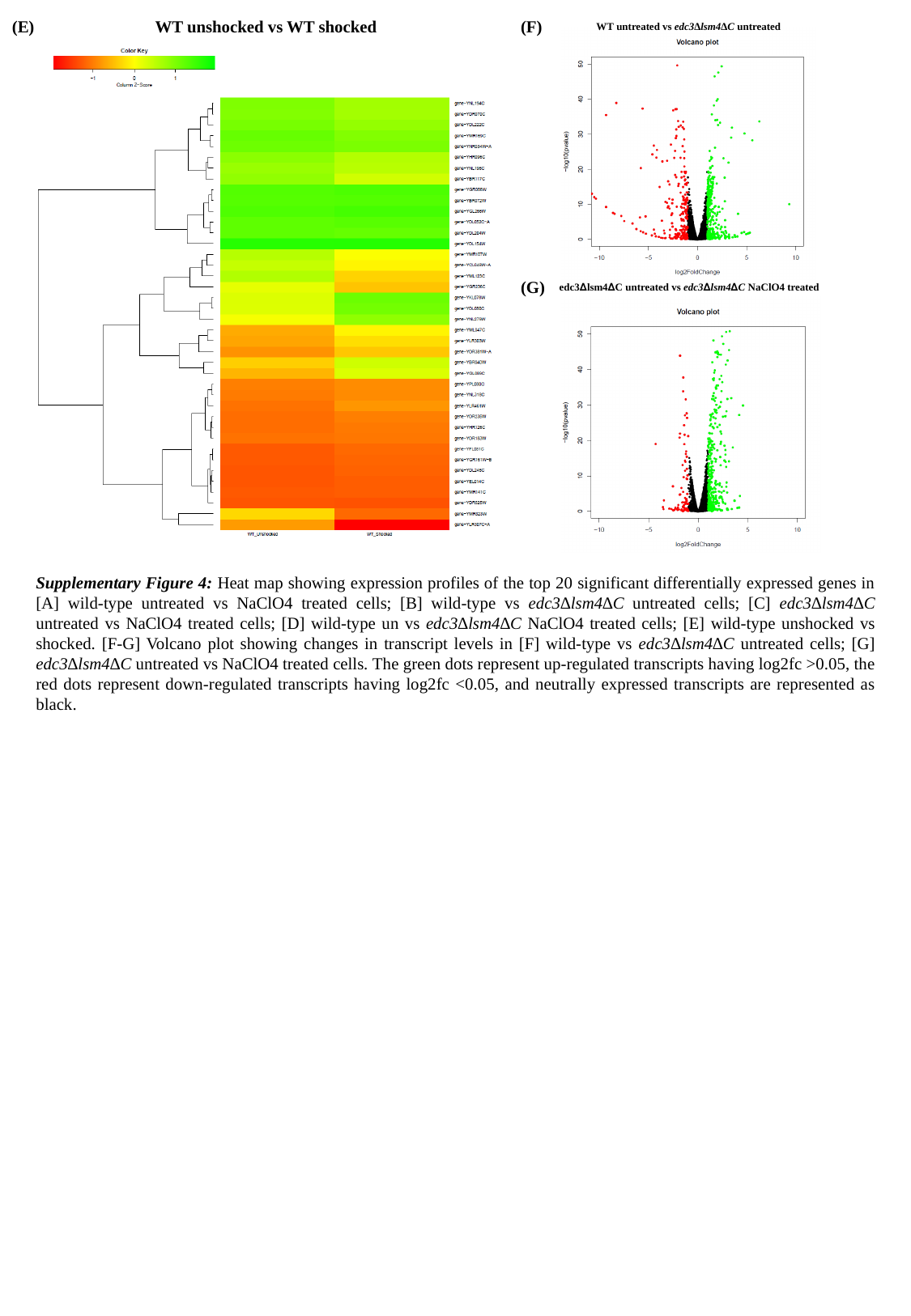

WT unshocked vs WT shocked
(F)
(E)
WT untreated vs edc3∆lsm4∆C untreated
(G)
 edc3∆lsm4∆C untreated vs edc3∆lsm4∆C NaClO4 treated
Supplementary Figure 4: Heat map showing expression profiles of the top 20 significant differentially expressed genes in [A] wild-type untreated vs NaClO4 treated cells; [B] wild-type vs edc3∆lsm4∆C untreated cells; [C] edc3∆lsm4∆C untreated vs NaClO4 treated cells; [D] wild-type un vs edc3∆lsm4∆C NaClO4 treated cells; [E] wild-type unshocked vs shocked. [F-G] Volcano plot showing changes in transcript levels in [F] wild-type vs edc3∆lsm4∆C untreated cells; [G] edc3∆lsm4∆C untreated vs NaClO4 treated cells. The green dots represent up-regulated transcripts having log2fc >0.05, the red dots represent down-regulated transcripts having log2fc <0.05, and neutrally expressed transcripts are represented as black.

### Slide 7
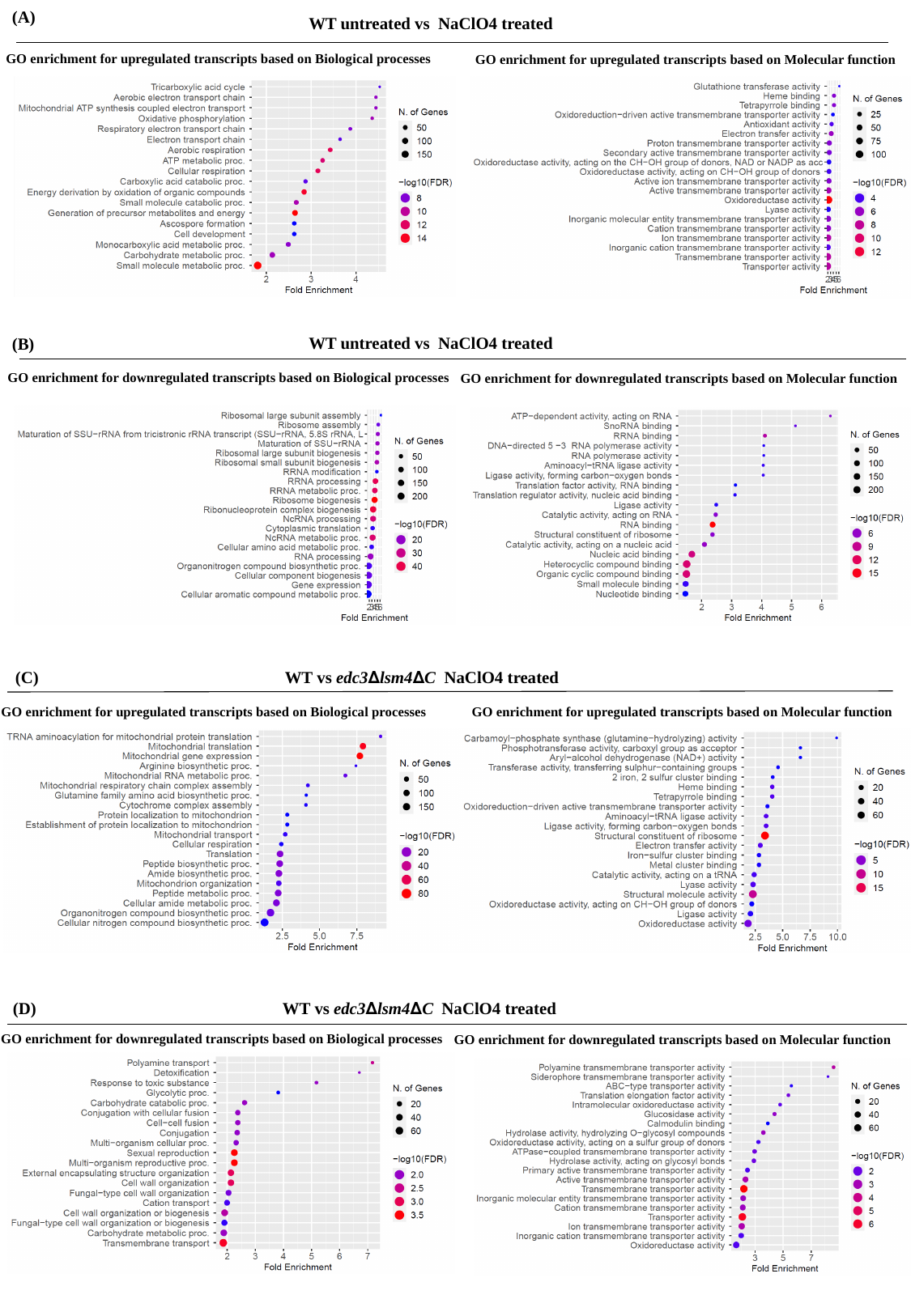

(A)
WT untreated vs NaClO4 treated
GO enrichment for upregulated transcripts based on Biological processes
GO enrichment for upregulated transcripts based on Molecular function
WT untreated vs NaClO4 treated
(B)
GO enrichment for downregulated transcripts based on Biological processes
GO enrichment for downregulated transcripts based on Molecular function
(C)
WT vs edc3∆lsm4∆C NaClO4 treated
GO enrichment for upregulated transcripts based on Biological processes
GO enrichment for upregulated transcripts based on Molecular function
(D)
WT vs edc3∆lsm4∆C NaClO4 treated
GO enrichment for downregulated transcripts based on Biological processes
GO enrichment for downregulated transcripts based on Molecular function

### Slide 8
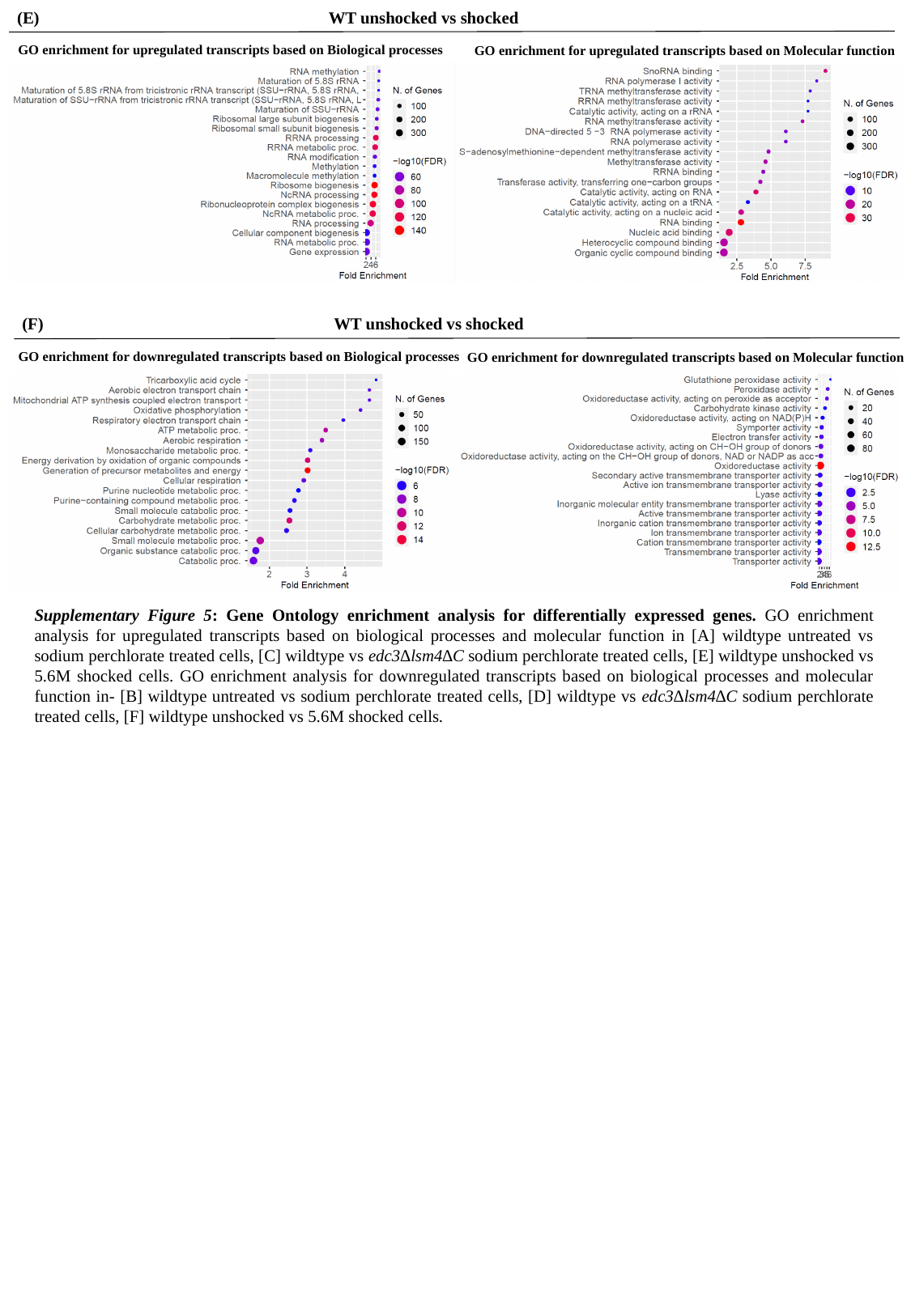

(E)
WT unshocked vs shocked
GO enrichment for upregulated transcripts based on Biological processes
GO enrichment for upregulated transcripts based on Molecular function
(F)
WT unshocked vs shocked
GO enrichment for downregulated transcripts based on Biological processes
GO enrichment for downregulated transcripts based on Molecular function
Supplementary Figure 5: Gene Ontology enrichment analysis for differentially expressed genes. GO enrichment analysis for upregulated transcripts based on biological processes and molecular function in [A] wildtype untreated vs sodium perchlorate treated cells, [C] wildtype vs edc3∆lsm4∆C sodium perchlorate treated cells, [E] wildtype unshocked vs 5.6M shocked cells. GO enrichment analysis for downregulated transcripts based on biological processes and molecular function in- [B] wildtype untreated vs sodium perchlorate treated cells, [D] wildtype vs edc3∆lsm4∆C sodium perchlorate treated cells, [F] wildtype unshocked vs 5.6M shocked cells.

### Slide 9
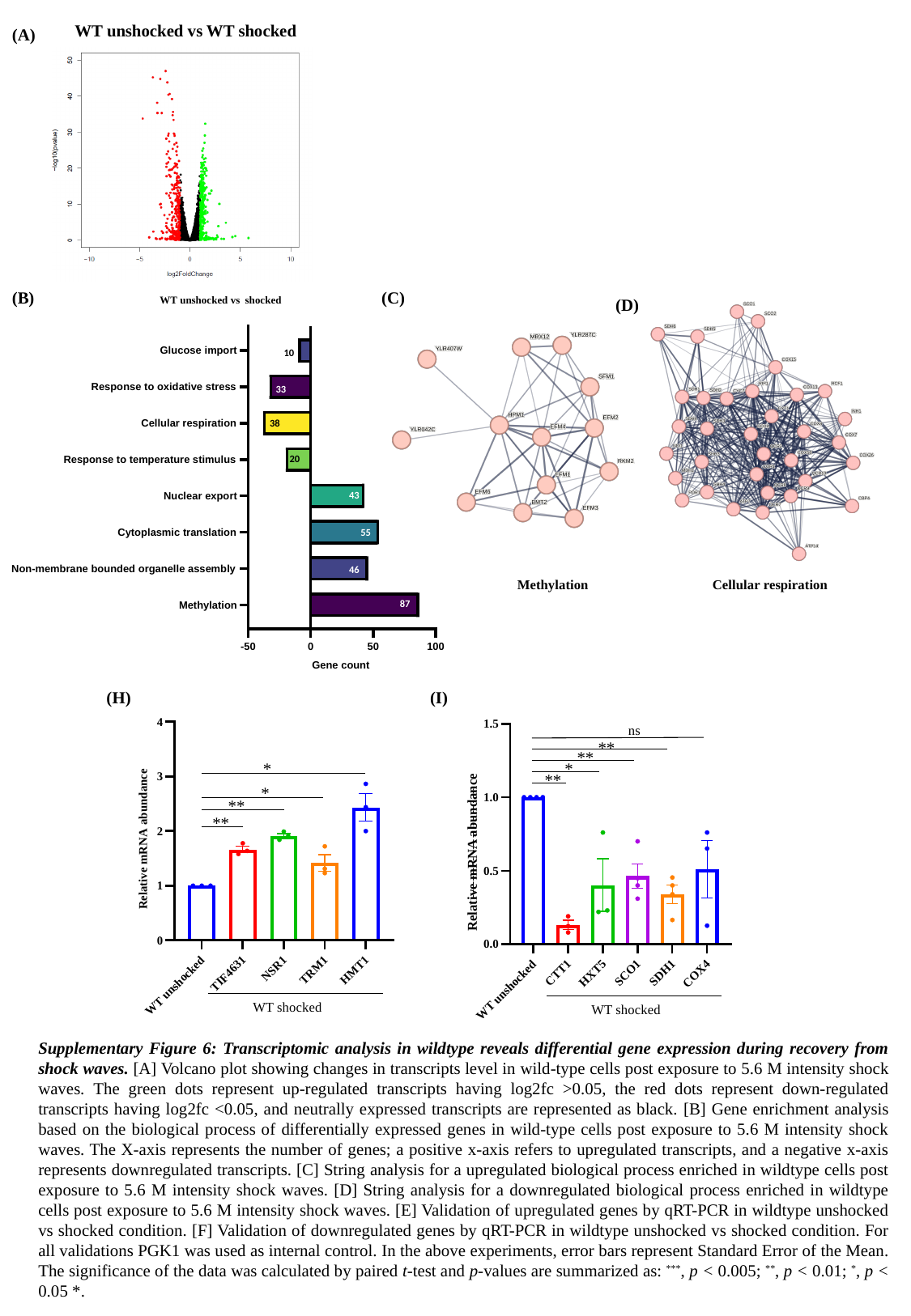

WT unshocked vs WT shocked
(A)
(B)
(C)
WT unshocked vs shocked
(D)
10
33
38
20
43
55
46
87
Methylation
Cellular respiration
(H)
(I)
ns
*
*
**
**
WT shocked
**
**
*
**
Relative mRNA abundance
WT shocked
Supplementary Figure 6: Transcriptomic analysis in wildtype reveals differential gene expression during recovery from shock waves. [A] Volcano plot showing changes in transcripts level in wild-type cells post exposure to 5.6 M intensity shock waves. The green dots represent up-regulated transcripts having log2fc >0.05, the red dots represent down-regulated transcripts having log2fc <0.05, and neutrally expressed transcripts are represented as black. [B] Gene enrichment analysis based on the biological process of differentially expressed genes in wild-type cells post exposure to 5.6 M intensity shock waves. The X-axis represents the number of genes; a positive x-axis refers to upregulated transcripts, and a negative x-axis represents downregulated transcripts. [C] String analysis for a upregulated biological process enriched in wildtype cells post exposure to 5.6 M intensity shock waves. [D] String analysis for a downregulated biological process enriched in wildtype cells post exposure to 5.6 M intensity shock waves. [E] Validation of upregulated genes by qRT-PCR in wildtype unshocked vs shocked condition. [F] Validation of downregulated genes by qRT-PCR in wildtype unshocked vs shocked condition. For all validations PGK1 was used as internal control. In the above experiments, error bars represent Standard Error of the Mean. The significance of the data was calculated by paired t-test and p-values are summarized as: ^***^, p < 0.005; ^**^, p < 0.01; ^*^, p < 0.05 *.

### Slide 10
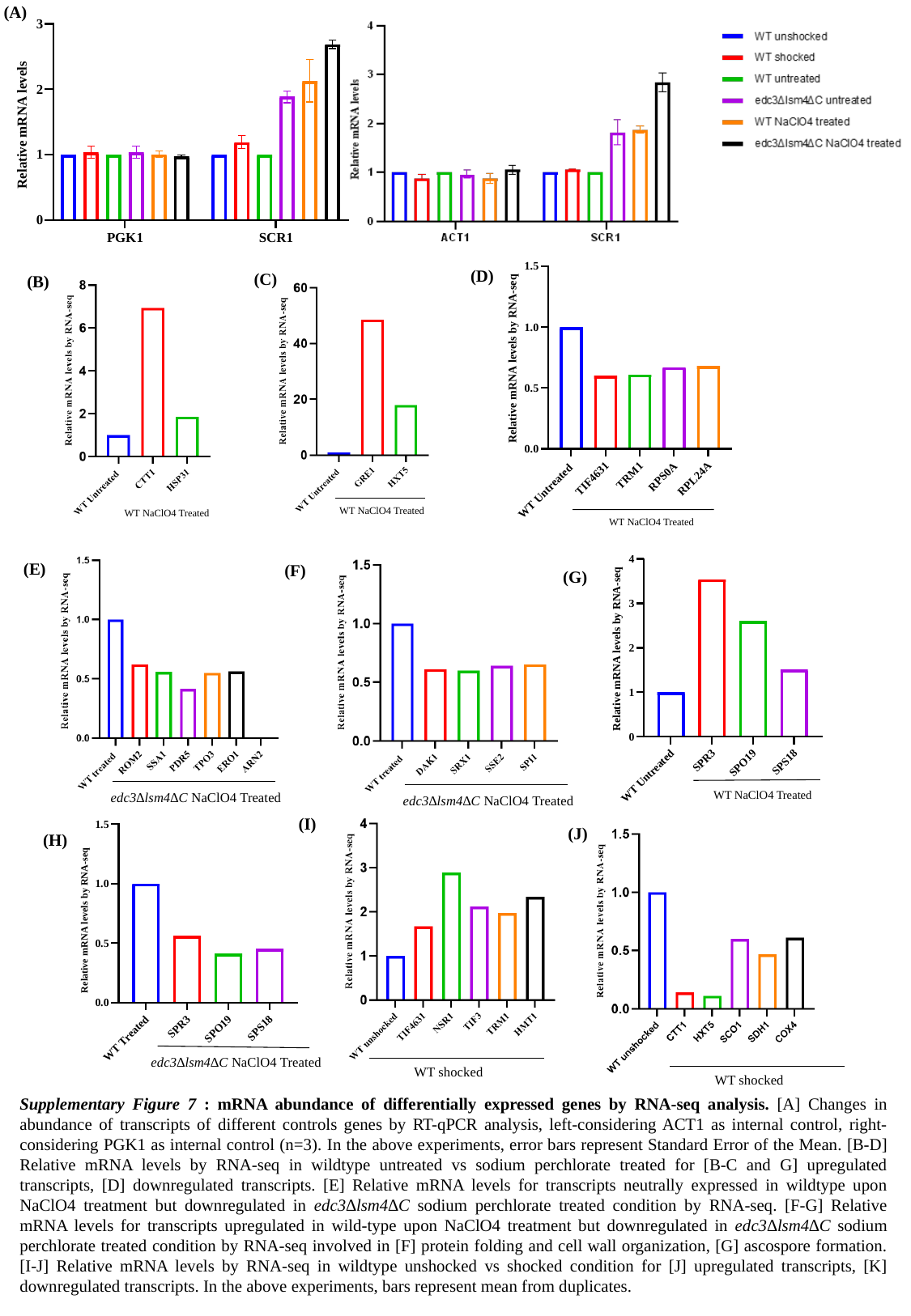

(A)
(D)
(C)
(B)
WT NaClO4 Treated
WT NaClO4 Treated
WT NaClO4 Treated
edc3∆lsm4∆C NaClO4 Treated
(E)
(F)
(G)
WT NaClO4 Treated
edc3∆lsm4∆C NaClO4 Treated
(I)
(J)
(H)
edc3∆lsm4∆C NaClO4 Treated
WT shocked
WT shocked
Supplementary Figure 7 : mRNA abundance of differentially expressed genes by RNA-seq analysis. [A] Changes in abundance of transcripts of different controls genes by RT-qPCR analysis, left-considering ACT1 as internal control, right-considering PGK1 as internal control (n=3). In the above experiments, error bars represent Standard Error of the Mean. [B-D] Relative mRNA levels by RNA-seq in wildtype untreated vs sodium perchlorate treated for [B-C and G] upregulated transcripts, [D] downregulated transcripts. [E] Relative mRNA levels for transcripts neutrally expressed in wildtype upon NaClO4 treatment but downregulated in edc3∆lsm4∆C sodium perchlorate treated condition by RNA-seq. [F-G] Relative mRNA levels for transcripts upregulated in wild-type upon NaClO4 treatment but downregulated in edc3∆lsm4∆C sodium perchlorate treated condition by RNA-seq involved in [F] protein folding and cell wall organization, [G] ascospore formation. [I-J] Relative mRNA levels by RNA-seq in wildtype unshocked vs shocked condition for [J] upregulated transcripts, [K] downregulated transcripts. In the above experiments, bars represent mean from duplicates.
